# The rise and fall of novel ecological communities

**DOI:** 10.1101/2020.06.02.131037

**Authors:** John M. Pandolfi, Timothy L. Staples, Wolfgang Kiessling

**Author notes:** Joint first authors (John M. Pandolfi, Timothy L. Staples).

## Abstract

Local and global environmental change is transforming ecological assemblages into new configurations, resulting in ecosystems with novel communities. Here we develop a robust methodology for the identification of novel communities, examine patterns in their natural chance of occurrence, and quantify the probability of local extinction, emigration, local origination and immigration in transitions to and from novel communities. Using a global dataset of Cenozoic marine plankton communities, we found the probability of local extinction, origination and emigration during transitions to a novel community increased up to four times that of background community changes, with the probability of species loss about equal to that of species gain. Although rare, once a novel community state emerged, the chance of shifting into another novel community state was five times greater than expected by chance. Thus, for marine planktonic communities at 100K year time scales, novel communities are particularly sensitive to further extinctions and community shift.

**One Sentence Summary:** Once developed, novel ecological communities face increased susceptibility to further shifts in species composition, with heightened extinction risk.

## Main text

Profound changes in the biodiversity of global ecosystems (*1, 2*) are leading to the formation of novel communities, where species composition and diversity are transformed into new, non-historical configurations (*3–5*). Key factors driving novel community emergence include the rapid pace of global climate change (*6–8*), breakdown of biogeographic barriers, species invasions and habitat degradation (*9–11*). These factors result in novel environments, new species combinations, and altered ecosystem functions (*12*). Little research has focused on the demographic drivers both during the transition to novel communities, and in subsequent time periods after they develop. Here we introduce a reproducible, objective, and quantitative approach to detect novel communities from time series compositional data, apply this approach to investigate the frequency and transition probabilities of community novelty over long temporal scales, and explore the demographic processes that influence the transition during, and after, the rise of novel community states.

The conceptual basis of novel ecosystems spans multiple disciplines, including the linked social-ecological systems of conservation biology, the no-analog communities of plant paleoecology and the emerging eco-climatic literature on novel climates. For conservation biologists, novel ecosystems contain historically unprecedented combinations of species, often with altered ecological functions that are driven by human agency, including the way in which human values interact with those ecosystems (*13*). The no-analog community concept from plant paleoecology focuses on the taxonomic composition or environmental framework of past communities that are compositionally unlike any found today (*14*). A variety of quantitative analytical approaches have been used to understand how and why past (*15–17*) or even future (*4*) vegetation dynamics in focal ecosystems differ from the present. Early emphasis on the role of novel climates in ecological change (*14, 18*) has prompted more recent robust quantitative investigation of the geographic variation and temporal distribution of novel climates (*7*), including under various Representative Concentration Pathway (RCP) scenarios (*19, 20*).

We build on these previous approaches to provide a novelty detection framework that can be applied to any taxonomic group from any ecosystem from any time period or timescale, and enables comparative ecosystem approaches to understand general trends, causes and consequences of novel communities on a global scale.

### A new framework for evaluating novelty in ecological communities

Our framework is based on three conceptual pillars, which provide the foundation for detecting novelty in ecological communities from compositional data within time series. First, our framework quantifies two separate criteria of ecological novelty. Second, we assess novel communities in their temporal context. Third, our approach makes assessments of novelty within each time series, so spatial variation in taxonomic composition is not considered for the detection of ecological novelty.

#### Two criteria for ecological novelty

We have used an early concept of novelty from the modern ecological literature where novel ecosystems are defined as those that are ‘rapidly being transformed into new, non-historical configurations’ (*3*). This definition excludes the prerequisite that novel communities must be driven by human activities, and frames novelty solely in ecological terms. It also decomposes novel communities into two components: (1) faster compositional turnover than is normally observed in the system, and (2) a shift to a composition that is substantially different from any historical state. Novel communities occur where both criteria are met. Notably, different types of novelty also occur when communities satisfy only one criterion, creating two additional categories of novelty. Quantifying this definition allows us to detect and compare multiple aspects of ecological novelty.

#### Temporal position

Most existing methods for quantifying novelty are largely based on spatial (or spatiotemporal) datasets and are not designed for making space-restrictive assessments, or working with time series data in which temporal order matters. However, the temporal positioning of communities is critical to evaluate their novelty components against historical backgrounds. We assessed novel communities within long time series, with their relative temporal position preserved, so they are assessed relative to past, but not future, ecological compositions. Our framework incorporates the notion that species composition is always in flux, and that a singular ‘historical baseline’ does not exist.

The dissimilarity in composition between two adjacent time bins within a time series quantifies the pace of community change (i.e. ‘rate-of-change’). We termed this dissimilarity ‘instantaneous dissimilarity’ (Fig 1A). We quantified the shift to an unprecedented composition as the smallest dissimilarity between a target community and all previous states (similar to *4*), treating communities from earlier in the time series as a collection of ‘previously observed community states’. We termed these dissimilarities ‘cumulative dissimilarity’ (Fig 1A). A small cumulative dissimilarity suggests that the composition is similar to one that has been previously observed; a large cumulative dissimilarity indicates that the target composition is very different from previous states.

**Figure 1.**
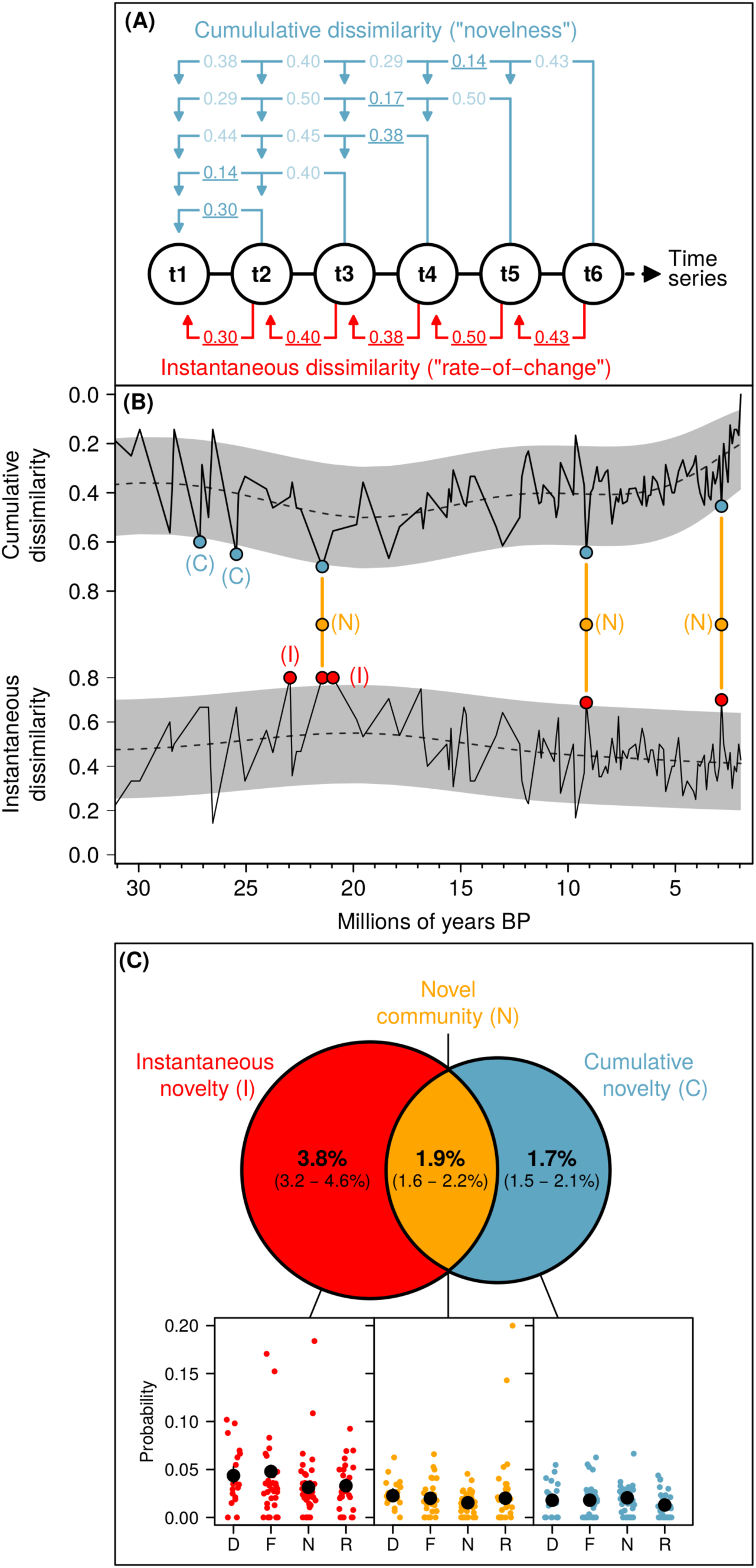
The novel community-detection framework and application to marine plankton. **(A)** Method for calculating instantaneous (red) and cumulative dissimilarity (blue) in time series compositional data. Instantaneous dissimilarities are calculated between pairs of adjacent time series samples (“bins”). Cumulative dissimilarities are the smallest dissimilarity in community composition between a target bin and all previous bins (shown as <u>underlined</u> values). (**B)** Example of our novel detection framework in a single time series. Black lines are the observed instantaneous and cumulative dissimilarities calculated for 100,000-year sampling bins along a single time series. Dashed line is the mean expected dissimilarity obtained from two generalized additive models. The grey shading indicates the upper and lower 95% predictive boundary. Bins that exceeded each boundary were classified as either ‘instantaneous novelty’ (I, in red), or ‘cumulative novelty’ (C, in blue). Bins classified as both were considered to be true novel communities (N, in orange). Note that the y-axis on the cumulative dissimilarity plot is reversed: values increase from top to bottom. (**C)** The Venn diagram shows the estimated probability of a community being classified, over all four marine planktonic groups. Colored points in subplots show proportion of time series bins categorized as each novelty classification. Black points are mean probabilities for each planktonic group (D = diatoms, F = foraminifers, N = calcareous nannoplankton and R = radiolarians), with 95% confidence intervals as error bars (most error bars are obscured behind mean estimates).

#### Within-time series assessments and expectations

We consider novelty to be a space-restricted attribute, where a community state could still be novel even when it was observed in other time series, so long as it was unprecedented in that community’s history. Given that community composition is always in flux from stochastic and deterministic processes, our within-time series approach let us define novelty not just as high dissimilarity, but higher dissimilarity than expected for that community state, capturing time series trends in instantaneous and cumulative dissimilarity (Fig. 1B). We generated expectations for both criteria of novelty along each time series using parametric spline-based models. These splines were free to increase or decrease through time to reflect changing expectations while retaining the time-ordering of the compositional data. Rather than focusing on a global threshold, we applied locally weighted thresholds for detecting novel communities. We quantified novelty firstly by comparing observed dissimilarity scores to expected dissimilarity distributions generated by these models. We then classified outliers that exceeded probabilistic thresholds in these distributions as novel communities. This gave us a pool of classified novel communities that could be aggregated across time series and used in further analyses. Some aspects of this approach are mirrored in the eco-climatic literature, which converted no-analog climatic comparisons into standardized anomalies of reference climate variation (*19*). This research differs from our approach by using short-term environmental variability to standardize cross-space comparisons, whereas our approach standardizes and makes comparisons strictly through time for each time series.

Our framework quantifying novelty results in four categories of compositional change: (1) a faster than expected shift in composition (‘instantaneous novelty’ - I), (2) a community state more different to any past state than expected (‘cumulative novelty’ - C), (3) both instantaneous and cumulative novelty (‘novel community’ - N) (Fig. 1B), and (4) neither instantaneous nor cumulative novelty (‘background community’ - B).

## Results

We used our novelty-detection framework to investigate the global occurrence probability of community novelty in the Cenozoic marine plankton record using a global dataset of microfossil data from deep sea drilling cores. Our data were derived from the Neptune Sandbox (NSB) microfossil database (*21, 22*) (http://www.nsb-mfn-berlin.de/). Incorporating modernized taxonomy and age models allowed us to build community data for groups of taxa across geochronological time (Fig. 2). Each time series in our analyses is a single or multiple cores from sites grouped into Longhurst biogeographical provinces (*23*), with species presence and absence grouped separately for four groups of marine plankton (calcareous nannoplankton, foraminifers, radiolarians and diatoms) every 100,000 years over the last 65 m.y. (Fig. 2). To evaluate the robustness of our results, we conducted additional comparative analyses, with novelty assessed at individual core locations (figs. S1-3), with low richness communities excluded (figs. S4-6), with varying sampling bin widths (200 ka – 500 ka: figs. S7-9) and excluding potential reworking (‘Pacman’ analysis: figs. S10-12). Results from all of these additional comparative analyses were similar to those presented below.

**Figure 2.**
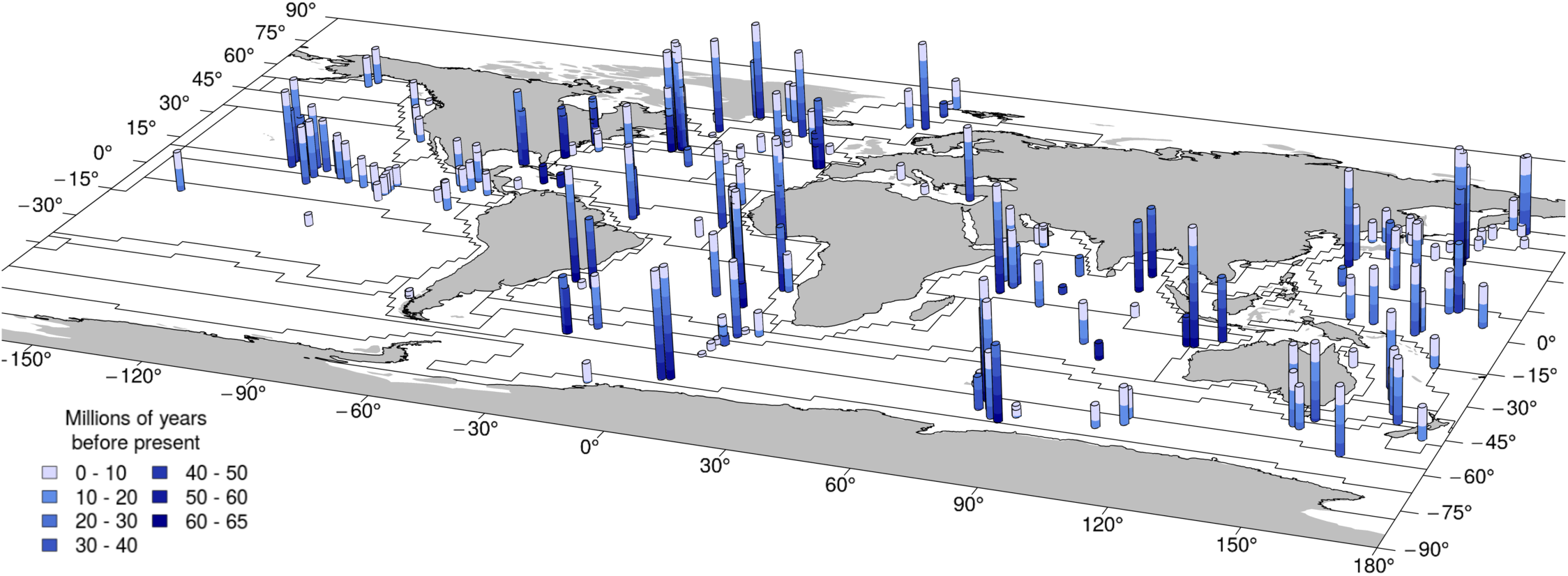
Map of Neptune Sandbox core locations, lengths and age ranges. Cylinders represent a single core, or the aggregate of multiple cores taken at a single location (i.e. one ‘site’). Cylinder length correlates with the duration of the time series at a particular site, broken into 10 million year bands. Band color corresponds to the age of that segment. Some sites contained community data for more than one planktonic group (calcareous nannoplankton, foraminifers, radiolarians and diatoms), but this detail is not shown here. Black polygons show the boundaries of Longhurst biogeographical provinces. Cores were aggregated into Longhurst provinces for analysis.

### Probability of novelty emergence and community state transitions

The probability of novel community emergence was consistent across the four marine planktonic groups (1.6 to 2.2%), with the probability of instantaneous novelty ranging from 3.2 to 4.6%, and cumulative novelty ranging from 1.5 to 2.1% (Fig. 1C). These probabilities provide an estimate for the frequency that novel planktonic communities arose under natural marine conditions in the absence of human impacts. Probabilities were sensitive to alpha threshold (width of predictive boundary in novel detection framework), which was set to 5% (Fig 3A). However, the proportional overlap among the three classifications was largely stable (Fig. 3B, C), and there was a strong positive Pearson’s correlation between instantaneous and cumulative dissimilarity scores (r = 0.77). Thus, subsequent analyses built on particular sets of novel communities are unlikely to be an artifact of our alpha threshold.

**Figure 3.**
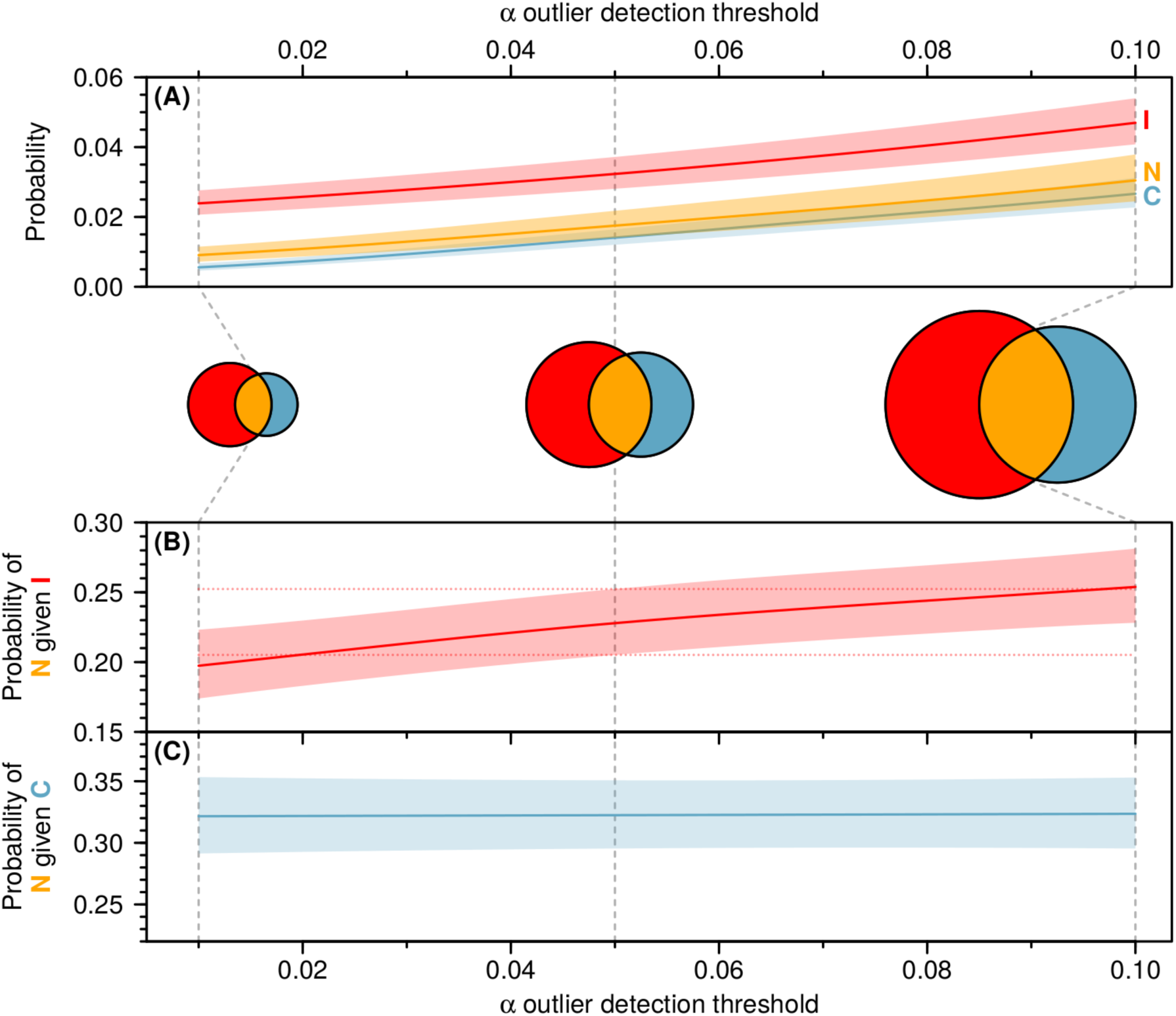
Sensitivity of novel emergence probabilities to alpha outlier detection threshold. **(A)** Models of probability of novelty emergence based on the cut-off threshold used to generate prediction intervals in our novelty detection framework. Lines are mean predicted probability, with colored polygons representing 95% confidence intervals. **(B)** and **(C)** show modeled probability of a novel community occurring given one type of ecological novelty. These are equivalent to the proportional overlap of the red circle with the blue circle in the central Venn diagrams, and vice versa, as the cut-off threshold becomes less stringent. The Venn diagrams are descriptive, highlighting the result that while the probability of detecting novelty increases as cut-off threshold becomes less stringent, the proportional overlap of novelty remains relatively consistent. I = Instantaneous novelty (red), C = Cumulative novelty (blue), and N = Novel community (orange).

We tested for signals of temporal autocorrelation of community classifications through time, examining whether transitions from one novel community to another were more likely to occur than by chance. To achieve this, we estimated the observed probabilities of pairwise transitions from one state to another (n = 16). We then compared these observed probabilities with non-autocorrelated expectations for each transition: the occurrence probabilities of each state, multiplied together.

Most observed transitions were from background to background communities (Fig. 4). Transitions between the three types of novelty and a background state were much rarer (‘Transitions to and from novelty’ in Fig. 4), and comparable to the emergence probabilities of novelty (reported above in Fig. 1C). Some of these transitions also occurred significantly less often than expected by chance. Transitions from one novelty state to another were rarer still, and, with the exception of one transition, led to subsequent novelty between two and ten times more often than expected (‘Transitions between novelty categories’ in Fig. 4). Thus, despite the infrequent occurrence of novelty throughout our time series, transitions between novelty states were disproportionately followed by subsequent novelty states, demonstrating a heightened propensity for novelty to beget further novelty.

**Figure 4.**
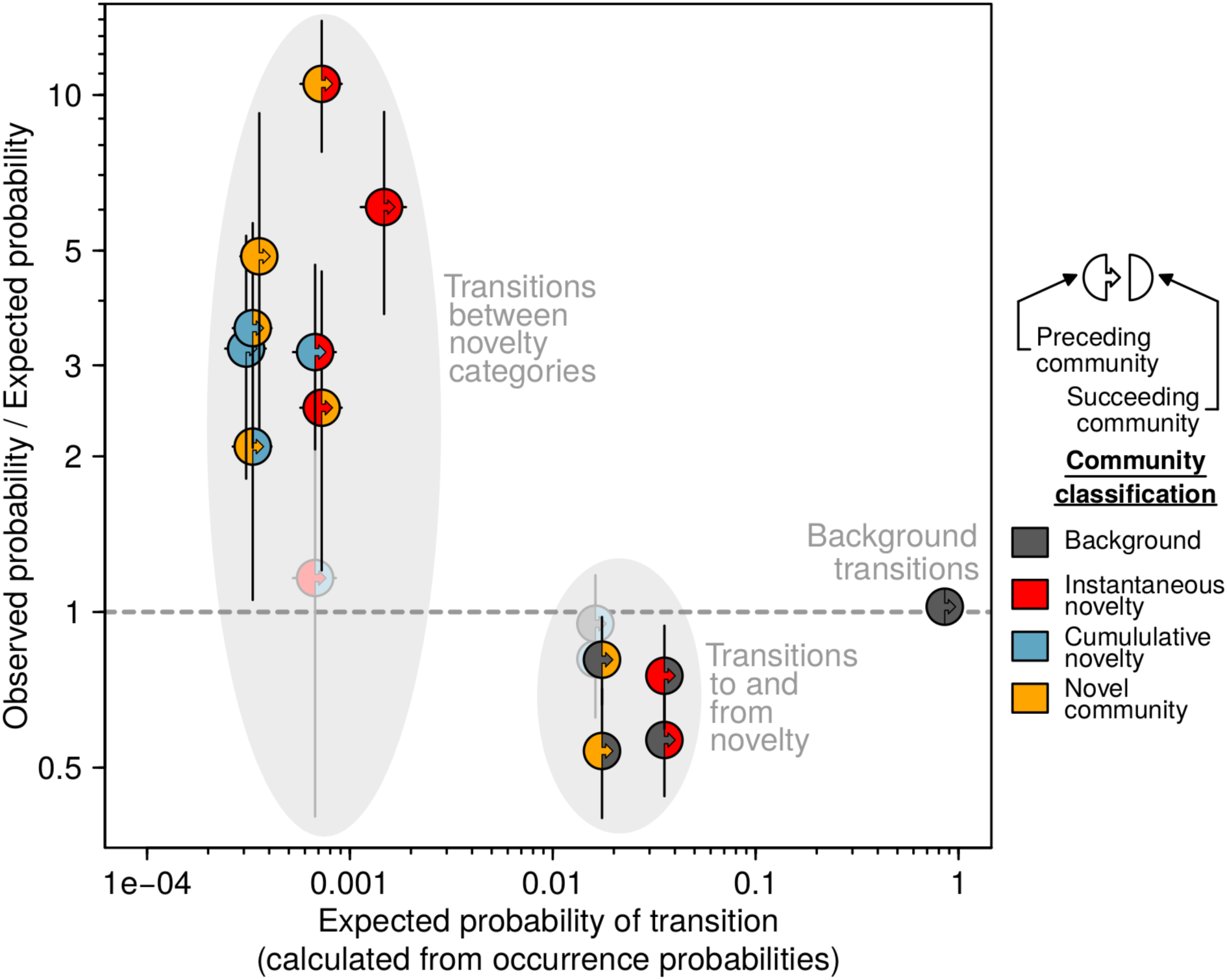
Expected probability of transitions between each of our four community classifications, and the ratio of observed transition probabilities to expected probabilities (table S5). Expected transition probabilities were estimated by multiplying the probabilities of occurrence for the preceding and succeeding classification from occurrence models (those used in the Venn diagram of Fig. 1C). Observed transition probabilities were estimated using binomial generalized linear mixed-effects models. Points are halved and dual-colored; the left-hand and right-hand color represent the classification of the preceding and succeeding community, respectively. Y-axis points are ratios: values greater than one indicate transition probability was higher in observed data than expected, and vice versa. Colored points are transitions where observed to expected ratio was significantly different to one: faded points had confidence intervals that crossed one. Note that both axes are ln-transformed.

### Demographic drivers of novelty

We evaluated the role of four demographic processes in driving transitions between one community state and another. ‘Local extinction’ (the permanent loss of a species from a time series) and ‘emigration’ (a transient loss with the species reappearing later in the time series) lead to species loss, whereas ‘local origination’ (the first occurrence of a species in a time series), and ‘immigration’ (a species subsequently reappearing after a temporary disappearance) lead to species gain. Background transitions act as a reference baseline for comparing demographic probabilities. We examined both unique loss or gain events (local extinction or origination) versus temporary loss (emigration) or re-colonizing events (immigration) and summed taxonomic gain vs taxonomic loss.

Species turnover was a major feature in the transition to novelty, with transitions to instantaneous novelty and background communities showing a small trend towards greater taxonomic gain and transitions to cumulative novelty and true novel communities showing small trends towards greater taxonomic loss (Fig. 5A). On average, we found taxa in the preceding community had a 10-14% probability of going locally extinct in the transition to a novel community. This is more than twice as high as transitions to background communities (local extinction, Fig. 5B). Similarly, taxa present in the subsequent novel community had a 18-27% probability on average of being entirely new to the time series (local origination, Fig. 5C), more than four times as high as transitions to background communities. Species gain in the transition to instantaneous novelty (I) was driven more by species recolonization (immigration, Fig. 5C) than by the origination of new taxa, which is consistent with instantaneous novelty being a large shift to a state similar to past states. In contrast, species gain in transitions to new, previously unseen community states (C) and novel communities (N) came about through the addition of new taxa, through invasion or evolution.

**Figure 5.**
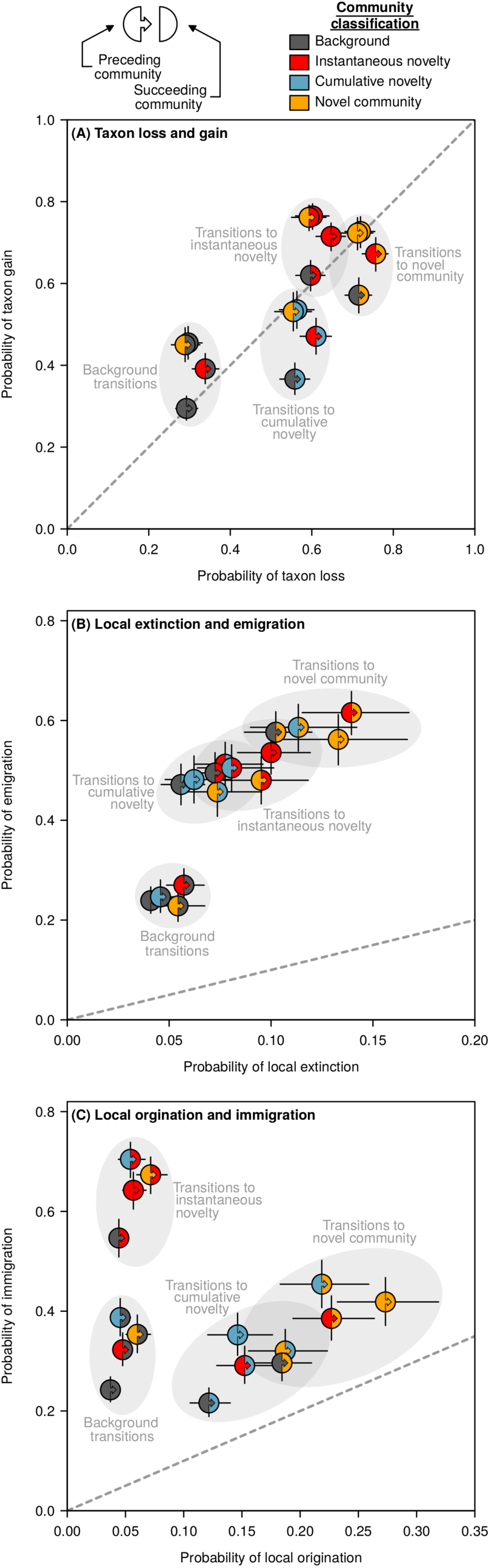
Mean probabilities of taxonomic loss or gain in the transition between two communities along a time series. Points are halved and dual-colored; the left-hand and right-hand color represents the classification of the preceding and succeeding community, respectively. (**A**) Total taxonomic loss plotted against total taxonomic gain. (**B**) Taxonomic loss plotted as the probability of local extinction (taxon disappears from time series and does not return) against the probability of emigration (species disappears transiently). (**C**) Taxonomic gain plotted as the probability of local origination (taxon appears in time series for the first time) against the probability of immigration (subsequent re-emergence of a previously-present taxon). The dashed line shows a 1:1 ratio and bars are 95% confidence intervals. Predictions were obtained from generalized linear mixed-effects models (model summaries shown in tables S6-S11).

In contrast to the transition *to* novelty states, taxonomic turnover played a relatively minor role in the subsequent transition *from* novelty (I, C or N) to background communities (Fig. 5A). This suggests there was no heightened extinction risk after a novel community emerged. However, when novel communities transition to a subsequent, different, novel state, these transitions were accompanied by increased probabilities of demographic changes, including local extinction (Fig. 5B) and local origination (Fig. 5C). Pairing this result with the increased risk of subsequent novel communities following other novel communities (Fig. 2), naturally-occurring novel communities recorded from the deep sea marine record have the potential to cascade through multiple novelty transitions, each associated with elevated rates of extinction and origination.

### Variation among the marine planktonic groups

Analysis of individual marine planktonic groups shows consistent patterns for expected probabilities of community state transition, particularly with background transitions, and transitions to and from novelty (Fig. 6). Transitions between novel community states were also similar, except that the calcareous nannoplankton showed a much higher (up to 20x higher than expected) propensity for novel communities to transition to instantaneous novelty than the other three planktonic groups. Patterns for demographic drivers are mostly consistent among taxa and with the overall trends (Fig. 7). However, the calcareous nannoplankton show the least difference in taxonomic turnover between background transitions and those to any type of novelty (Fig. 7). Moreover, planktonic foraminifera show lower probabilities of extinction than radiolarians and diatoms, and the diatoms show lower probabilities of origination than radiolarians and foraminifera in the transition to true novel communities (Fig. 7). We also found differences among taxa in the temporal distribution of novelty emergence. For example, novel community emergence peaked for the radiolarians and foraminifera between 55 and 54 MYA, and for foraminifera and calcareous nannoplankton between 30 and 23 MYA (fig. S13). In contrast, diatoms showed no such trends in novelty community emergence through time. These spikes in novelty community emergence appear to be associated with warming trends (fig. S13). Future work is needed to examine the spatial congruence in the occurrence of novelty among the four taxa, as well as a more detailed analysis of both the environmental drivers of novelty through time and the degree to which demographic drivers of novelty are influenced by accelerated rates of climate and environmental change. Although patterns within the planktonic groups generally follow the overall trends in the data, differences point to the utility of our framework in differentiating trends in novelty and their demographic drivers among taxa.

**Figure 6.**
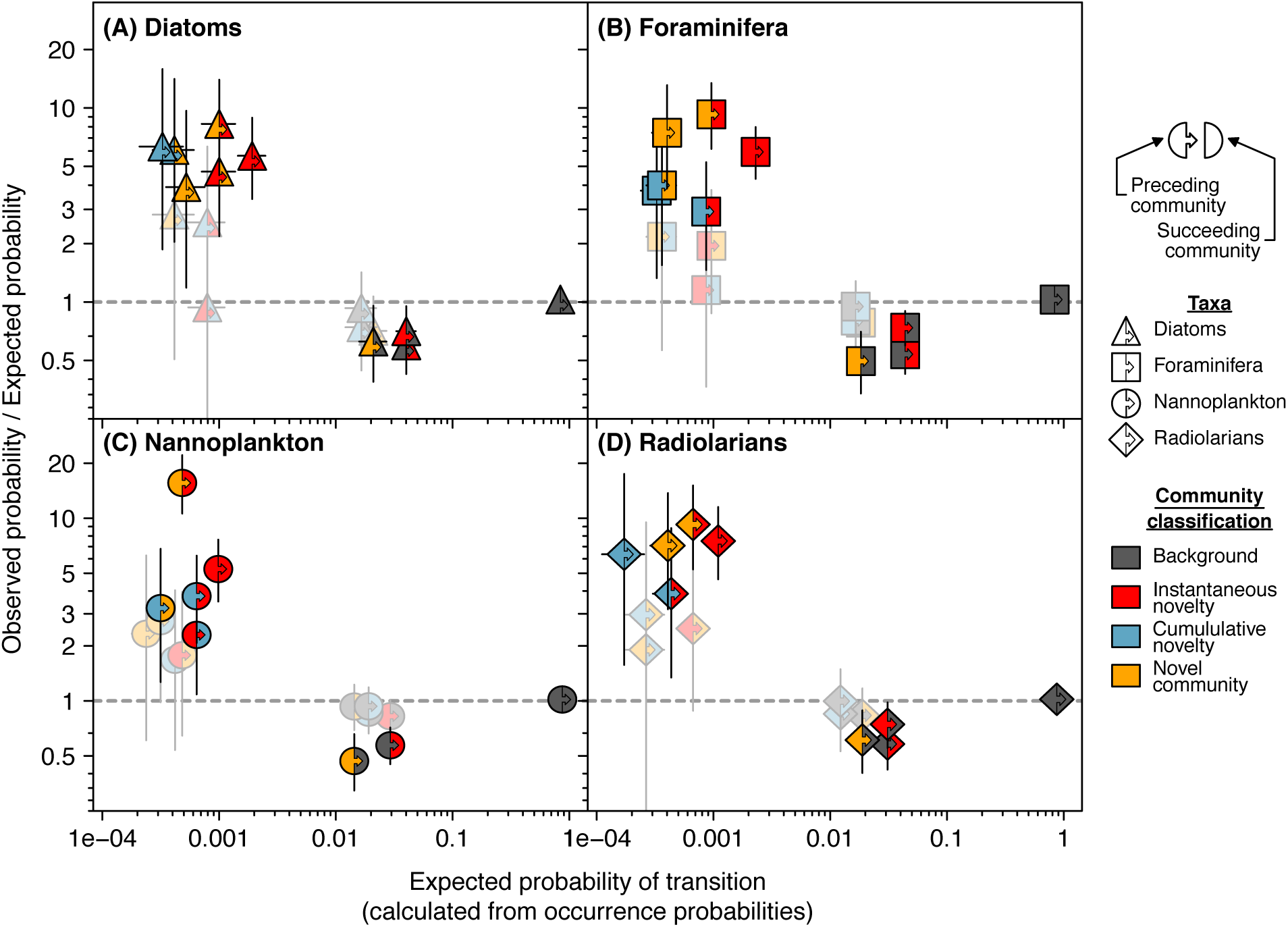
Expected probability of transitions between each of our four community classifications, and the ratio of observed transition probabilities to expected probabilities, modeled separately for each group of planktonic taxa. See Fig. 4 caption for further explanation.

**Figure 7.**
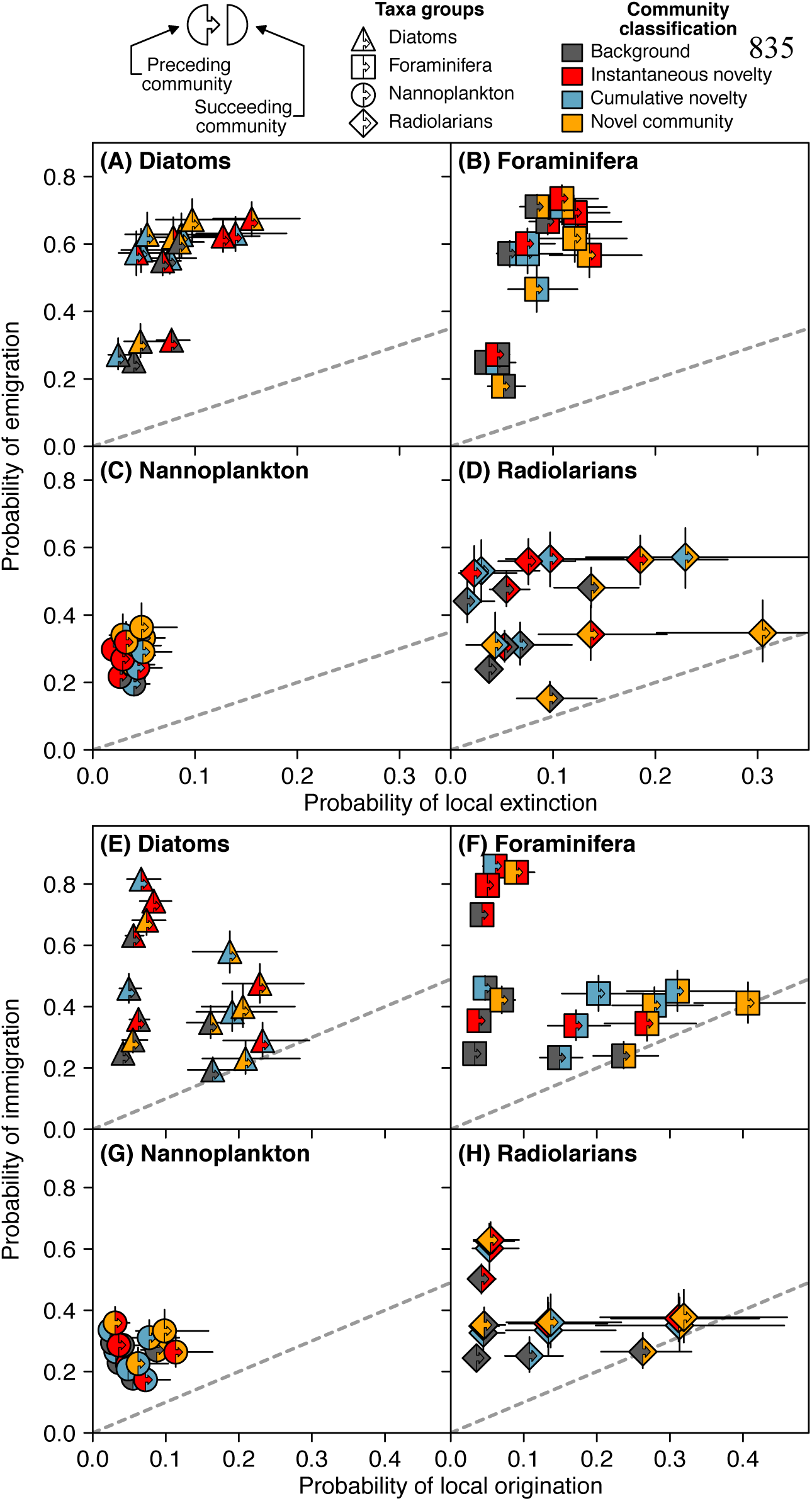
Mean probabilities of taxonomic loss or gain in the transition between two communities along a time series, with separate estimates for each group of planktonic taxa. **A-D** correspond to Fig 5B, and **E-H** correspond to Fig 5C. See Fig. 5 caption for further explanation.

### Confronting potential bias

#### Detection errors

Detection errors are not restricted to data gathered from the fossil record, but they are worth mentioning in the context of large databases that have been amassed from a number of different published data sources. Because the goal of different publications from which NSB is derived ranged from biostratigraphy to biodiversity (*24*), some of the published data was not intended to document biodiversity, with some workers discarding a significant proportion of the species within their samples, especially in the Radiolaria (*25*). However, we found no systematic differences in overall patterns between the Radiolaria and the other three groups of taxa that would suggest any sampling bias.

#### Taphonomic and sampling bias

Due to the constant and favorable preservation conditions, the deep-sea microfossil record is exceptionally rich and complete. The absence of any systematic increase or decrease in novelty over time (fig. S13), renders long-term preservational biases unlikely. However, coring sites might still be subject to reworking, age-model errors and other taphonomic uncertainties. To account for these, we ran a set of subsequent analyses on the NSB data with the oldest 5% and youngest 5% of species occurrences excluded using the ‘Pacman’ method (*26*). The results from this analysis were virtually indistinguishable from the analyses including all species occurrences (fig. S10-12).

The ‘Pacman’procedure does not correct for the bias of incomplete sampling within the stratigraphic ranges of species. Heterogeneous sampling completeness among aggregated Longhurst provinces may have affected our ability to take the metrics of species turnover at face value. We used locally weighted sampling probability to assess the variation of sampling completeness among Longhurst provinces and plankton groups and excluded Longhurst provinces with low sample completeness prior to statistical analyses (*27*).

## Discussion

Our study focused on four marine plankton groups with very different life-history attributes, including autotrophic and heterotrophic taxa, as well as different skeletal mineralogy (calcareous vs siliceous). Yet the probability of novelty, observed occurrence of community transitions and patterns in demographic drivers of novelty were all largely consistent across the four taxonomic groups (Figs. 6, 7). This remarkable similarity suggests general ecological constraints that govern the rise and fall of novelty in these planktonic communities.

Our approach provides a quantitative tool to examine the frequency and drivers of novelty in ecological communities that can be broadly applied at any temporal scale for any taxonomic group, and any ecosystem. It is particularly useful for understanding beta diversity in modern ecosystems, for the comparative analysis of the frequency, persistence and drivers of community change and novelty in human-dominated vs past ecosystems, and for synthetic macroecological analyses across ecosystem and taxonomic boundaries. However, we caution that additional factors must be considered. For example, long-term studies that connect ecological and evolutionary time scales need to consider the role of origination (and extinction) in ecosystems from the deep past when comparing the frequency of novelty with present ecosystems. Our analyses of novelty in marine plankton communities over 65 million years inevitably includes an evolutionary component that could lead to higher estimates of novelty in fossil communities relative to modern communities. On the other hand, short-term fluctuations in community structure would be averaged out over the 100 ka time bins analyzed in the fossil time-series data, potentially leading to lower estimates of past novelty than in modern communities sampled over much shorter time intervals (*28*). Generation times also vary widely across the biosphere and these must be considered alongside the temporal duration of the study for any comparative analyses of the frequency or drivers of community novelty. These and other factors will need to be assessed before the analysis of novelty in fossil assemblages can be used as an ecological baseline for the frequency of novelty in living ecosystems. Nevertheless, our analyses provide a foundation for understanding the past frequency and demographic drivers of novelty in natural settings over an extended time frame and in the absence of human impacts.

It is estimated that more than 35% of modern terrestrial communities now exist in a novel state (*29*), albeit these estimates and the criteria used for novelty are not generally based on quantitative or comparative analyses. Of particular concern to conservation biologists is the fate of modern communities that have transitioned into novel states: will these new configurations of species persist or are they likely to lead to further community change or even ecosystem collapse? Novelty in our fossil marine planktonic communities most often transitioned to a background state, implying that many shifts to novel states may persist through time. Moreover, changes in species composition *per se* may not necessarily reduce the functioning of an ecosystem where functional redundancy exists. However, deep time transitions to novel communities in the marine plankton exhibited both species loss and species gain, but ultimately resulted in shifts in community composition with at least twice the probability of local extinction. We also observed that novel communities were five times more likely to give rise to other novel communities than expected. While there are fundamental differences in ecological and environmental factors governing species composition over different time scales, our results from fossil communities raise the intriguing hypothesis that modern novel ecological communities could also be more likely than expected to experience transitions to subsequent novel community states, driven in part by further extinctions. Our novelty framework is well suited to further study of modern time series to test this directly.

Although we are aware of the dangers in direct comparison of ecological dynamics over vastly different time scales, our results raise the possibility that efforts to reduce extinction risk are consistent with the active management and conservation of novelty in modern ecological communities (*30–32*). Under the influence of human impacts, failing to improve the conditions that brought about the transition to novelty may facilitate further novelty accompanied by additional species extinction. Our results contravene any notion that a fixed historical baseline of a community can or should be the only conservation goal for novel communities (*3*). Rather, ecosystem management should also focus on how to prevent transitions to additional previously unseen ecosystem states that are linked with heightened extinction risk.

## Materials and methods

### Experimental Design

All of our data were retrieved from the Neptune Sandbox Berlin (*21*) (NSB, http://www.nsb-mfn-berlin.de/), a global dataset of marine microfossil occurrence data collected from deep-sea cores of international ocean drilling projects (Fig. 2). We collated Cenozoic NSB entries from 249 unique time series for four sets of planktonic groups: calcareous nannoplankton (171 time series), foraminifers (165 time series), radiolarians (147 time series) and diatoms (109 time series). These data were downloaded from the NSB data portal on the 10^th^ October 2019.

For each group of taxa, we treated synonymized taxonomic IDs provided by NSB as ‘species’. Relative abundance data were only available for a very small proportion of NSB time series, so we chose to use presence-absence data. Despite this decision, our novelty framework is equally applicable for abundance data. We aggregated the presence of species in sites into the same Longhurst province (Fig. 2) (*23*), separately for each of our four taxonomic groups. Each Longhurst province is thus treated as a time series for each plankton group.

The time between sampling events varied, both within and among time series. Therefore, we aggregated compositional samples along each times-series into standardized time windows (herein ‘bin’), representing a single ‘community’. We binned our time series to a constant sampling resolution of 100,000 years (see figs. S7-9 for analyses at larger bin widths) due to the uncertainties inherent in modeling age over geologic time. At this temporal grain, we likely only detected very large and prolonged shifts in community composition, and missed short, ephemeral shifts that occurred on shorter time-scales. The maximum sampling bin age was 65 million years, yielding 651 potential bins per time series. We excluded time series from Longhurst provinces with insufficient sampling, which we defined as those that either contained fewer than ten 100,000-year sampling bins, or the presence of fewer than ten distinct taxa across the time series.

Heterogeneous sampling completeness among aggregated Longhurst provinces may have affected our ability to take the metrics of species turnover at face value. Sampling completeness can be evaluated with the simple concept of sampling events/sampling opportunities. We used locally weighted sampling probability to assess the variation of sampling completeness among Longhurst provinces and plankton groups (*27*). This metric measures the number of taxa detected in three consecutive bins (three-timers, ^3^t) and the number of taxa detected before and after a focal bin but not within it (part timers, ^P^t). The overall sampling probability in Longhurst provinces is thus given by ^3^t/(^3^t + ^P^t). Sampling completeness was similar between taxonomic groups but varied among Longhurst provinces. We removed provinces with sampling completeness below the 5% quantile for each group as potentially taphonomically biased (fig. S14). These sites were excluded from further analyses. Our novel framework and subsequent statistical analyses were performed in R version 3.6.2 (*33*).

### Identifying novelty in ecological time series

#### Novel framework methodology

We quantified two criteria of ecological novelty: (1) faster than expected compositional turnover, (2) to a state that is more different to any historical state than expected. To do so, we extracted the dissimilarity between each community and its predecessor as a metric of ‘rate-of-change’, which we termed ‘instantaneous dissimilarity’. We also extracted the smallest dissimilarity between a community and all previous sampling bins, to act as a metric of ‘novelness’, which we termed ‘cumulative dissimilarity’ (Fig. 1A). We estimated dissimilarities between each pair of communities in each time series using the Jaccard index (vegdist function, vegan package (*34*)). Both Jaccard and Bray-Curtis exclude joint unshared species, increase linearly with ecological distance, and are linear until distances are large (*35*). We chose the Jaccard index as it is a metric transformation of Bray-Curtis (*36*). Jaccard is commonly used in ecological studies because it quantifies overlap in the species actually observed in the two communities, does not weight unshared species, and cuts out shared zeroes. When we employed a number of different metrics, they all tended to identify similar proportions of novel communities, and identified many of the same communities in the NSB dataset (fig. S15).

We tested whether the instantaneous and cumulative dissimilarity of a community exceeded expectations generated using two generalized additive models (GAMs). We considered it unlikely that dissimilarities would change in a linear manner over time, so we chose to fit spline-based models that are free to take varying shapes dependent on the data. We used the GAMs to estimate 95% predictive boundaries through time separately for instantaneous and cumulative dissimilarity. We opted to use GAMs instead of other spline-based options using running-mean smoothers such as LOESS, as GAMs have robust penalization methods that prevent over-fitting (*37*). Since many dissimilarity indices are bound between zero and one (including Jaccard), we fit the GAMs with beta errors and logit link function (transforming dissimilarities equal to zero and one as Y = (y × (n − 1) + 0.5)/n, where *y* values were the raw dissimilarities and *n* was the number of bins in the time series (*38*). Each model contained a single fixed effect, “age”, which was the center of each sampling bin (e.g., sampling bin 0 – 100K years BP, “age” = 50 K). This age variable was fit as a cubic regression spline, with the spline complexity determined by restricted maximum likelihood. This reduces the risk of spline over-fitting, but is less prone to under-fitting than other penalization methods (such as generalized cross validation) (*39*). We set maximum spline complexity in the GAM formula to k = −1, the default in the GAM function, which automatically selects a maximum spline complexity. We also re-ran our detection framework with k = 10, which allows for strongly non-linear trends. This process gave identical results, albeit with much longer computation time.

The two GAMs estimated the mean predicted instantaneous and cumulative dissimilarity along the time series. Our definition of novel communities relies on detecting bins with dissimilarities that exceed these expectations. To identify these outliers, we established beta distributions at each bin along each time series using the beta distribution. The beta distribution function family in R (e.g., qbeta, pbeta) uses the α and β shape parameters to calculate the beta distribution shape. The beta GAM does not produce α and β directly, but instead provides µ (the predicted dissimilarity at each bin on the logit scale) and Φ (the precision parameter, a single value for the entire model). We converted these into α and β using α = µ × Φ and β = Φ − α (mgcv package (*37*)), and used these α and β values to calculate a beta distribution for each bin along the time series.

We compared each instantaneous and cumulative dissimilarity to the beta distribution of expectations using these shape parameters (pbeta function). This resulted in two *p*-values for each bin along the time series. These *p*-values standardize novelty onto continuous probability scales, with lower probabilities corresponding with more outlying (i.e. more “novel”) communities in a time series relative to expectations. These *p*-values are useful for comparing novelty in a manner that is conceptually similar to the methodology used by Mahony et al. (*19*) to standardize climatic conditions.

Our goal in this study was to identify a pool of communities exhibiting different types of novelty, which we used to identify unique patterns and signals of novelty. We chose to identify probability outliers, where observed dissimilarities exceeded the 95% predictive boundary of the beta distribution at that bin. We calculated 95% predictive boundaries as the 95^th^ quantile of the bin’s beta distribution (qbeta function). Bins with observed dissimilarities that exceeded these thresholds were classified as either ‘instantaneous novelty’ (I), based on the model of instantaneous dissimilarity, or ‘cumulative novelty’ (C), based on the model of cumulative dissimilarity. Bins that qualified as both instantaneous and cumulative novelty constituted a ‘novel community’ (N). Bins that qualified as neither type of novelty were classified as ‘background’ communities (B). This gave us four possible community classifications. Our approach in detecting cumulative novelty echoes that of Radeloff et al. (*4*) except that they make comparisons: 1) to a fixed baseline set, and 2) among time periods across space.

Our method of outlier detection applied a cut-off at alpha = 0.05. Prior work on cross-space novelty also compared continuous metrics of novelty to cut-off thresholds (*4, 16*). However, our predictive boundaries can vary not only between time points within each time series, but also at the same time point in different time series. This standardizes our detection of novelty in a manner that these previous works do not (but see *19* for a conceptually similar approach). This means our 95% predictive boundaries do not simply identify the top 5% of dissimilarities in the NSB data, or along each time series. Some time series had no instances of novelty, and others exhibited much more than 5% (Fig. 1C). Not only is our methodology for estimating expected dissimilarity thresholds flexible along the time series, it has two conservative components that reduce the risk of misidentifying novel communities: 1) blocks of time in a given time series with consistently high dissimilarities raise locally-weighted expectations, and 2) time series with high residual variation in dissimilarities have wider 95% prediction intervals (via larger Φ). Both of these components increase both the *p*-values of continuous novelty, and the predictive threshold to be detected as an outlier.

In our framework, a particular community state can only be classified as a ‘novel community’ (N) once in a given time series. This can be thought of as the ‘emergence’ of a novel community. If we observed the exact same composition again in the same time series, cumulative dissimilarity would be zero, and this repeated composition could not qualify as cumulative novelty (C). If the repeated composition was present in the bin after we observed a ‘novel community’, our framework would classify it as a ‘background’ community (B). If a previous ‘novel’ composition was observed again later in the time series, it could only be classified as either ‘background’ or ‘instantaneous novelty’ (I), depending on the magnitude of the immediate bin-to-bin change.

We tested whether our framework reliably identified novel communities using two simulations (See Supplementary Materials for details). In the first, we created synthetic ecological time series expressing particular signals of turnover, with *a priori* expectations for where novel communities occurred (figs. S19-20). In the second, we used a random walk simulation of species turnover to generate ecological time series to compare to observed probabilities from the NSB data (table S12).

#### Sensitivity of cumulative novelty test to edge effects

Given that our cumulative novelty test uses a pool of previous community states, it may be sensitive to edge effects. Early in a time series, the pool of previous states is small, which may make it more likely to obtain high cumulative dissimilarities. To test for edge-effects, we modeled whether outliers from our two dissimilarity GAMs were more likely early in time series. We used two binomial generalized linear mixed effects models (logit link function: glmer function, lme4 package (*40*)), one for instantaneous dissimilarity, and one for cumulative dissimilarity. We treated each bin within each time series as a trial, with the outliers in each of our framework models constituting a success, and non-outliers constituting a failure. Our predictor variable was the bin position from the start of the time series, treated as a continuous variable, and including a squared interaction effect to examine non-linear changes in probability as bin position increased. We fit random intercepts to each of our four planktonic groups, and then separately to each time series within these groups.

We found that the probability of detecting outliers was indeed substantially higher in the initial time bins, particularly for cumulative dissimilarity (fig. S16). We tested whether removing the initial time-points from each time series would correct this edge effect. We ran our novel detection framework on the entirety of each time series, then removed the first five time-bins and re-ran the models specified above. With these initial bins removed, relationships between time series position and novelty probability were essentially flat (fig. S16D-F). Accordingly, we used the dataset with the initial five time bins removed for all analyses.

We also found that novel communities were more likely to occur in communities with fewer species (fig. S17). As we were unable to determine whether these were true ecological signals or a symptom of incomplete sampling, we also ran our novelty detection framework and analyses excluding communities with fewer than five species (fig. S4-6). Results were nearly identical to those that included all communities.

#### Sensitivity of novel framework to prediction interval width

We chose a 95% predictive interval to define novelty, which equates to an alpha cutoff of 0.05. This is an almost universal statistical cutoff, although not without criticism (*41*). We investigated the influence of the alpha threshold on the probability of detecting outliers in our two dissimilarity tests, as well as the probability of classifying as instantaneous novelty, cumulative novelty or as a novel community (Fig. 3). We set alpha to multiple values, from 0.01 to 0.10 in units of 0.005, running our novel detection framework using each value. We then modeled the probability of identifying each of our three novelty classifications separately for each alpha value. The models used were binomial mixed-effect GAMs (GAMMs, gamm4 function, gamm4 package (*42*)). The response variables were sets of weighted trials, identical to those used to generate Fig. 1B (see Statistical analyses section below for details). We fit alpha threshold as a continuous fixed effect, set as a cubic regression spline to model changes in probability across tested alpha thresholds. We set nested random intercepts for each model, with individual Longhurst provinces nested within planktonic taxa groups.

We also quantified the proportional overlap of our two outlier tests using two additional GAMs. These models equate to testing the probability of detecting a novel community from the pool of instantaneous novelty communities, or the pool of cumulative novelty communities. For these models, we treated each time series as a set of weighted trials, with each novel community constituting a success. In these models, we treated the number of instantaneous or cumulative novelty communities in each time series as the number of trials, rather than the number of time-points along the time series.

As reported in the main text, we found that the overall probability of detecting outliers increased as we relaxed our alpha threshold, which is expected. The probability of overlap between our instantaneous and cumulative novelty groups, however, remained remarkably consistent regardless of cut-off. Even as the absolute number of outliers increased, the proportion of them that satisfied both novelty criteria to be classified as ‘novel communities’ was similar (Fig. 3).

### Statistical Analyses

Our novelty framework resulted in four pools of novel community classifications nested within individual Longhurst province time series, one for each group of planktonic taxa. We used these classifications in the following analyses: (1) we modeled the average occurrence probability of each community classification across all taxa, and within each of our four taxonomic groups; (2) we tested for patterns of temporal autocorrelation between one community classification and another; and (3) we quantified probabilities of taxon loss and gain between pairs of community classifications along each time series, breaking them into four demographic drivers.

#### The probability of emergence of novelty and novel communities

We calculated the occurrence probability using binomial regression models, for each of our four community classifications (Fig. 1B), and compared probabilities between the four planktonic groups. We used two sets of models, one set treating group as levels of a categorical predictor (“fixed-taxa models”: glm function), and one treating groups as levels of a random intercept (“random-taxa models”: glmer function, lme4 package (*40*)). The random-taxa models were intercept-only, and contained no fixed effects. In each model, we treated the bins in each time series as a set of trials, with each bin constituting a single success or failure. Success was defined as the occurrence of the target classification. All other community classifications were treated as failures. By treating each time series as a set of trials, the models weighted observed probabilities based on time series length, which prevented longer time series from unduly influencing model estimates. We included time series from all four planktonic groups in each model. The main results of these models (excluding the probability of a background classification) are summarized in the sub-panels of Fig. 1C.

#### Are observed probabilities of novelty transitions temporally auto-correlated compared with patterns expected by chance?

We estimated the ‘by chance’ or ‘expected’ probability of a given transition by multiplying the occurrence probabilities of the preceding and succeeding classifications together. As an example, the estimated probability of a transition from a novel community to instantaneous novelty occurring by chance is the occurrence probability of a novel community (0.019) multiplied by the occurrence probability of instantaneous novelty (0.038): an expected transition probability of 7.222e^−4^. Given four community classifications, there were 16 possible transitions from one classification to another.

We estimated the expected probability of each transition occurring using the random-taxa occurrence probability models (described above) from the previous analysis. We back-transformed estimates from a logit to probability scale (plogis function), and multiplied each pairwise combination of occurrence probabilities together. This meant that transitions to and from the same pair of states resulted in the same expected probability (e.g., probability of I → N was equivalent to the probability of N → I).

We estimated observed transition probabilities using GLMMs similar to those used above to estimate the occurrence probability, except we treated adjacent pairs of communities in each times-series as a single trial (e.g., bins one and two constituted one trial, bins two and three another etc.), with all pairs in a time series treated as a set of trials. Successes were pairs of bins that matched a particular transition, and failures were all other classifications. This resulted in 16 GLMMs, each modeling the probability of occurrence for one transition (“transition probability models”).

Contrasting observation with expectation allowed us to estimate effect size. We estimated effect sizes as the ratio of observed to expected transition probability. These effect sizes were multiplicative, where ratios of one indicated observed transition probabilities that were similar to expected probabilities. Ratios greater than one indicated that the transition was more likely to occur in the observed data than expected, and ratios less than one suggested the transition was less likely to occur than expected (Fig. 4).

Both the expected and observed transition probabilities were derived from models, and these probabilities contain uncertainty. We accounted for uncertainty by treating the modeled probabilities as Gaussian distributions. For the expected transition probabilities, we drew 100,000 occurrence probabilities for the preceding and succeeding classification from the random-taxa models. We treated the mean model estimate as the center of a Gaussian distribution, with a standard deviation equal to the standard error of the model estimate on the link-scale (logit). Once drawn, we back-transformed these probabilities onto the probability scale (plogis function). We then multiplied each set of preceding and succeeding classification probabilities together to give a distribution of expected transition probabilities. We repeated this process for each of the 16 possible transitions. These distributions were Gaussian, so we treated the mean of each distribution as the expected transition probability, and the 2.5% and 97.5% quantiles were used as upper and lower confidence intervals.

For the observed transitions probabilities, we drew 100,000 random estimates from the model coefficients of the transition probability models, and back-transformed these to the probability scale. We used these observed transition probability distributions along with the expected transition probability distributions to calculate our effect sizes. We did this by dividing each of our 100,000 observed transition probabilities by the corresponding entry in the expected transition probability distribution. This gave us 16 effect size distributions, which were also Gaussian. We used the mean of each effect size distribution as the effect size, with 2.5% and 97.5% quantiles as 95% confidence intervals.

#### Demographic drivers of novelty

We estimated whether the transition to a novel community, or a community detected as instantaneous or cumulative novelty, was associated with the rates of species gain or loss. We did this by modeling the probability of four categories of species turnover based on the classification of the preceding and succeeding community, and whether these differed from a background to background transition. The four categories of species turnover included taxonomic loss (local extinction and emigration) and taxonomic gain (local origination and immigration).

We used four GLMMs, one for each of our turnover categories (“turnover models”). All response variables were binary, where each taxon represented a success or failure from a set of trials, based on the number of taxa present in the preceding or succeeding bin. For the models using taxon loss, we set the number of trials to the number of taxa present in the preceding bin (i.e. the number of taxa that could be lost in the move to the succeeding bin). For the models using taxon gain, the number of trials was the number of taxa present in the succeeding bin (i.e. the number of taxa that could have appeared in the move to the succeeding bin).

These models contained nested random intercepts for each marine planktonic group, and Longhurst provinces within these groups. We used four fixed effects. The first two were preceding community classification and succeeding community classification, which were additive. The next two were continuous fixed effects. The first continuous fixed effect was the time lag between sampling bins. Not all possible bins were sampled in each time series. Larger gaps between samples may have resulted in greater species turnover, and we wanted to account for differences in turnover probabilities due to this time lag. We included the time difference between the preceding and succeeding bin as a continuous fixed effect to account for this time lag. After modeling, we estimated the probabilities for each transition category at the mean lag between bins.

The second continuous fixed effect was number of bins to the start and end of each time series, included only in the local origination and local extinction models, respectively. The probability of a species locally originating or becoming locally extinct is related to where the first and last occurrence, respectively, occurs in the time series. For example, local extinction was much more likely at the end of each time series, where a taxon has to be absent for fewer sampling bins to be classified as locally extinct. To detect the magnitude of this bias, we modeled the probability of local extinction and origination as a function of bins from the time series end and start, respectively, using GLMMs. The relationship between bin position (‘bins from time series start’ for local origination and immigration, and ‘bins from time series end’ for local extinction and emigration) and probability of local extinction and origination was exponential and decreased rapidly from bins one to ten (fig. S18).

To account for this edge effect in our species turnover models, we removed the first ten bins of each time series for the local origination and immigration models (as immigration count was linked to local origination count), and the last ten bins of each time series for the local extinction and emigration models. The removal of the ten bins at the start of each time series for the local origination model is in addition to the five we removed previously due to uncertainty of the cumulative novelty test (fig. S16).

Even after removing these bins, the probability of local extinction and origination was still negatively correlated with bin position (fig. S18). This residual relationship was linear, so we included bin position as a covariate in the models. Modeled turnover probabilities for each transition category were estimated at the mean bin position.

## Acknowledgments

This work was funded by the Australian Research Council’s Centre of Excellence for Coral Reef Studies (CE140100020) and by the Deutsche Forschungsgemeinschaft (DFG, German Research Foundation, KI 806/16-1). Data used in this study are available from the Neptune Sandbox microfossil database (http://www.nsb-mfn-berlin.de/index). R code to conduct all analyses and produce all tables and figures are available at Zenodo (URL) and Figshare (URL). We are grateful for discussion about this work with Stephen Jackson, David Lazarus, Jonathan Levine, and Richard Norris. The ideas for the paper were conceived by JMP and WK and enhanced by TLS; TLS undertook all analyses and developed analytical approaches with input from JMP and WK; JMP and TLS wrote the paper with WK contributing to writing and editing.

## Notes

### Competing Interest Statement

The authors have declared no competing interest.

## REFERENCES AND NOTES

1. M. Dornelas, N. J. Gotelli, B. McGill, H. Shimadzu, F. Moyes, C. Sievers, A. E. Magurran, Assemblage time series reveal biodiversity change but not systematic loss. Science. 344, 296–299 (2014).

2. S. A. Blowes, S. R. Supp, L. H. Antão, A. Bates, H. Bruelheide, J. M. Chase, F. Moyes, A. Magurran, B. McGill, I. H. Myers-Smith, M. Winter, A. D. Bjorkman, D. E. Bowler, J. E. K. Byrnes, A. Gonzalez, J. Hines, F. Isbell, H. P. Jones, L. M. Navarro, P. L. Thompson, M. Vellend, C. Waldock, M. Dornelas, The geography of biodiversity change in marine and terrestrial assemblages. Science. 366, 339–345 (2019).

3. R. J. Hobbs, E. Higgs, J. A. Harris, Novel ecosystems: implications for conservation and restoration. Trends Ecol. Evol. 24, 599–605 (2009).

4. V. C. Radeloff, J. W. Williams, B. L. Bateman, K. D. Burke, S. K. Carter, E. S. Childress, K. J. Cromwell, C. Gratton, A. O. Hasley, B. M. Kraemer, A. W. Latzka, E. Marin-Spiotta, C. D. Meine, S. E. Munoz, T. M. Neeson, A. M. Pidgeon, A. R. Rissman, R. J. Rivera, L. M. Szymanski, J. Usinowicz, The rise of novelty in ecosystems. Ecol. Appl. 25, 2051–2068 (2015).

5. A. E. Magurran, How ecosystems change. Science. 351, 448–449 (2016).

6. D. Stralberg, D. Jongsomjit, C. A. Howell, M. A. Snyder, J. D. Alexander, J. A. Wiens, T. L. Root, Re-shuffling of species with climate disruption: a no-analog future for California birds? PLoS One. 4, e6825–e6825 (2009).

7. A. Ordonez, J. W. Williams, J. C. Svenning, Mapping climatic mechanisms likely to favour the emergence of novel communities. Nat. Clim. Chang. 6, 1104–1109 (2016).

8. B. R. Scheffers, L. De Meester, T. C. L. Bridge, A. A. Hoffmann, J. M. Pandolfi, R. T. Corlett, S. H. M. Butchart, P. Pearce-Kelly, K. M. Kovacs, D. Dudgeon, M. Pacifici, C. Rondinini, W. B. Foden, T. G. Martin, C. Mora, D. Bickford, J. E. M. Watson, The broad footprint of climate change from genes to biomes to people. Science. 354, aaf7671 (2016).

9. A. E. Lugo, T. A. Carlo, J. M. Wunderle Jr, Natural mixing of species: novel plant–animal communities on Caribbean Islands. Anim. Conserv. 15, 233–241 (2012).

10. A. Traveset, R. Heleno, S. Chamorro, P. Vargas, C. K. McMullen, R. Castro-Urgal, M. Nogales, H. W. Herrera, J. M. Olesen, Invaders of pollination networks in the Galapagos Islands: emergence of novel communities. Proceedings. Biol. Sci. 280, 20123040 (2013).

11. A. Anton, N. R. Geraldi, C. E. Lovelock, E. T. Apostolaki, S. Bennett, J. Cebrian, D. Krause-Jensen, N. Marbà, P. Martinetto, J. M. Pandolfi, J. Santana-Garcon, C. M. Duarte, Global ecological impacts of marine exotic species. Nat. Ecol. Evol. 3, 787–800 (2019).

12. R. J. Hobbs, L. E. Valentine, R. J. Standish, S. T. Jackson, Movers and Stayers: Novel Assemblages in Changing Environments. Trends Ecol. Evol. 33, 116–128 (2018).

13. R. J. Hobbs, E. S. Higgs, C. M. Hall, in Novel Ecosystems: Intervening in the New Ecological World Order (2013), Wiley Online Books, pp. 58–60.

14. J. W. Williams, S. T. Jackson, Novel climates, no-analog communities, and ecological surprises. Front. Ecol. Environ. 5, 475–482 (2007).

15. J. W. Williams, B. N. Shuman, T. Webb, Dissimilarity analyses of late-quaternary vegetation and climate in eastern North America. Ecology. 82, 3346–3362 (2001).

16. W. Finsinger, T. Giesecke, S. Brewer, M. Leydet, Emergence patterns of novelty in European vegetation assemblages over the past 15 000 years. Ecol. Lett. 20, 336–346 (2017).

17. V. Iglesias, C. Whitlock, T. R. Krause, R. G. Baker, Past vegetation dynamics in the Yellowstone region highlight the vulnerability of mountain systems to climate change. J. Biogeogr. 45, 1768–1780 (2018).

18. J. W. Williams, S. T. Jackson, J. E. Kutzbach, Projected distributions of novel and disappearing climates by 2100 AD. Proc. Natl. Acad. Sci. U. S. A. 104, 5738–5742 (2007).

19. C. R. Mahony, A. J. Cannon, T. Wang, S. N. Aitken, A closer look at novel climates: new methods and insights at continental to landscape scales. Glob. Chang. Biol. 23, 3934–3955 (2017).

20. M. C. Fitzpatrick, J. L. Blois, J. W. Williams, D. Nieto-Lugilde, K. C. Maguire, D. J. Lorenz, How will climate novelty influence ecological forecasts? Using the Quaternary to assess future reliability. Glob. Chang. Biol. 24, 3575–3586 (2018).

21. D. Lazarus, Neptune: A marine micropaleontology database. Math. Geol. 26, 817–832 (1994).

22. C. Spencer-Cervato, The Cenozoic deep sea microfossil record: explorations of the DSDP/ODP sample set using the Neptune database. Palaeontol. Electron. 2 (1999).

23. A. R. Longhurst, Ecological Geography of the Sea (Academic Press, San Diego, ed. 2., 2007).

24. J. Renaudie, D. Lazarus, P. Diver, NSB: an expanded and improved database of marine planktonic microfossil data and deep-sea stratigraphy. EarthArXiv (2019), doi: 10.31223/osf.io/97se5.

25. D. Lazarus, N. Suzuki, J.-P. Caulet, C. Nigrini, I. Goll, R. Goll, J. K. Dolven, P. Diver, A. Sanfilippo, An evaluated list of Cenozic-Recent radiolarian species names (Polycystinea), based on those used in the DSDP, ODP and IODP deep-sea drilling programs. Zootaxa. 3999, 301–333 (2015).

26. D. Lazarus, M. Weinkauf, P. Diver, Pacman profiling: a simple procedure to identify stratigraphic outliers in high-density deep-sea microfossil data. Paleobiology. 38, 144–161 (2012).

27. J. Alroy, Dynamics of origination and extinction in the marine fossil record. Proc. Natl. Acad. Sci. 105, 11536–11542 (2008).

28. D. B. Kemp, K. Eichenseer, W. Kiessling, Maximum rates of climate change are systematically underestimated in the geological record. Nat. Commun. 6, 8890 (2015).

29. E. Marris, Ecology: Ragamuffin earth. Nature. 460, 450–453 (2009).

30. A. M. Truitt, E. F. Granek, M. J. Duveneck, K. A. Goldsmith, M. P. Jordan, K. C. Yazzie, What is Novel About Novel Ecosystems: Managing Change in an Ever-Changing World. Environ. Manage. 55, 1217–1226 (2015).

31. S. J. Capon, G. J. Palmer, Turning over a new leaf: the role of novel riparian ecosystems in catchment management. Solut. J. 9 (2018).

32. S. Clement, R. J. Standish, Novel ecosystems: Governance and conservation in the age of the Anthropocene. J. Environ. Manage. 208, 36–45 (2018).

33. R Development Core Team, A Language and Environment for Statistical Computing. R Found. Stat. Comput. (2019), (available at http://www.r-project.org).

34. J. Oksanen, F. G. Blanchet, M. Friendly, R. Kindt, P. Legendre, D. McGlinn, P. R. Minchin, R. B. O’Hara, G. L. Simpson, P. Solymos, M. H. H. Stevens, E. Szoecs, H. Wagner, vegan: Community Ecology Package (2019), (available at https://cran.r-project.org/package=vegan).

35. D. P. Faith, P. R. Minchin, L. Belbin, Compositional dissimilarity as a robust measure of ecological distance. Vegetatio. 69, 57–68 (1987).

36. S. Kosub, A note on the triangle inequality for the Jaccard distance. Pattern Recognit. Lett. 120, 36–38 (2019).

37. S. N. Wood, Generalized additive models: An introduction with R, second edition (Chapman and Hall/CRC, ed. 2, 2017).

38. M. Smithson, J. Verkuilen, A better lemon squeezer? Maximum-likelihood regression with beta-distributed dependent variables. Psychol. Methods. 11, 54–71 (2006).

39. S. N. Wood, Fast stable restricted maximum likelihood and marginal likelihood estimation of semiparametric generalized linear models. J. R. Stat. Soc. Ser. B Stat. Methodol. 73, 3–36 (2011).

40. D. Bates, M. Mächler, B. M. Bolker, S. C. Walker, Fitting linear mixed-effects models using lme4. J. Stat. Softw. 67, 1–48 (2015).

41. D. J. Benjamin, J. O. Berger, M. Johannesson, B. A. Nosek, E.-J. Wagenmakers, R. Berk, K. A. Bollen, B. Brembs, L. Brown, C. Camerer, D. Cesarini, C. D. Chambers, M. Clyde, T. D. Cook, P. De Boeck, Z. Dienes, A. Dreber, K. Easwaran, C. Efferson, E. Fehr, F. Fidler, A. P. Field, M. Forster, E. I. George, R. Gonzalez, S. Goodman, E. Green, D. P. Green, A. G. Greenwald, J. D. Hadfield, L. V Hedges, L. Held, T. Hua Ho, H. Hoijtink, D. J. Hruschka, K. Imai, G. Imbens, J. P. A. Ioannidis, M. Jeon, J. H. Jones, M. Kirchler, D. Laibson, J. List, R. Little, A. Lupia, E. Machery, S. E. Maxwell, M. McCarthy, D. A. Moore, S. L. Morgan, M. Munafó, S. Nakagawa, B. Nyhan, T. H. Parker, L. Pericchi, M. Perugini, J. Rouder, J. Rousseau, V. Savalei, F. D. Schönbrodt, T. Sellke, B. Sinclair, D. Tingley, T. Van Zandt, S. Vazire, D. J. Watts, C. Winship, R. L. Wolpert, Y. Xie, C. Young, J. Zinman, V. E. Johnson, Redefine statistical significance. Nat. Hum. Behav. 2, 6–10 (2018).

42. S. Wood, F. Scheipl, gamm4: Generalized additive mixed models using mgcv and lme4 (2017), (available at https://cran.r-project.org/package=gamm4).

43. J. Zachos, M. Pagani, L. Sloan, E. Thomas, K. Billups, Trends, Rhythms, and Aberrations in Global Climate 65 Ma to Present. Science. 292, 686–693 (2001).

