## Supplemental Material for "The rise and fall of novel ecological communities"

Science AAAS

**This PDF file includes:**

Materials and Methods
Figs. S1 to S20
Tables S1 to S12

### 1 Supplemental Methods

#### 1.1 Novelty emergence probability and temperature change through time

We used Deep Sea $\delta^{18}O$ isotopic ratios (obtained from Zachos et. al. (*43*)) as a proxy for paleotemperature. We calculated mean $\delta^{18}O$ values for each 100K year sampling bin, and calculated the absolute change in $\delta^{18}O$ between each pair of bins to represent temperature change. We modeled direct correlation between temperature change and novel community emergence over time, and trends in both over time. Direct correlation was modeled using a binomial generalized linear mixed effects model, and included five fixed effects: temperature change, planktonic taxon group, their interaction, and two covariates used in models in the main text. These were the time lag between each time point and the previous, and the time point position along the time series. Trends in novel communities and temperature change were estimated using generalized additive models (GAMs). The novel community model fit years before present as cubic regression splines for each taxonomic group, with a maximum of 66 knots (one per million years in the study period), as well as lag between bins and bin position as per the correlation model in the main text. The temperature change GAM was fit the same, but only included the years before present spline term. Our time series derived from the Neptune dataset ranged over 65 million years where proxies for paleotemperature have been derived using Deep Sea $\delta^{18}O$ isotopic ratios (*43*). There was a weak positive relationship between temperature change and novel community emergence (Estimate = 2.397, SE = 1.40, $z$ = 1.71, $p$ = 0.09). The reason for the weak correlation is clearer when observing trends over time (fig. S13). Trends among taxa, and the change in novel community emergence based on temperature change, differed substantially through time. We found two periods of temperature change with corresponding spikes in novel community emergence in some taxa: 55-54 MYA for radiolarians and foraminifera, and 23-30 MYA for foraminifera and nannoplankton. Diatoms showed an almost flat probability rate of novel community emergence across the study period.

#### 1.2 Testing our framework using simulated communities

Given that our methodology is largely informed by the attributes of the time series, we tested the reliability of our framework in time series with particular types of compositional changes. Initially, we populated a 20 x 20 matrix to act as a template time series. Rows in the matrix were time points along the matrix, and columns were species. Each cell in the matrix (species abundance in a given time point) was seeded with an initial abundance of five, plus a random component estimated from a Poisson distribution ($\lambda$ = 1). These abundances were converted to relative abundances for each time point after any scenario-specific modifications (see below). We used this template matrix to construct various time series with patterns of compositional change, which we converted to dissimilarity matrices using the weighted Jaccard index (aka Ruzicka index). We note that these simulations use relative abundance composition data, while our analysis of the Neptune Sandbox is based on presence/absence data. Our novelty framework works via a dissimilarity matrix, regardless of data type. The top panels in figs. S19 and S20 show the dissimilarity from the base of each time series, which illustrate potential turnover patterns that are not exclusive to abundance or presence/absence data types.

##### 1.2.1 Stable state (fig. S19A, G, M)

This matrix represents a composition where species abundance only changes randomly around an equilibrium (where equilibrium is an abundance of five per species).

As expected, we did not find novel communities in this time series. This is evidence that even though our framework is informed by time series attributes, it is unlikely that noise around a stable state will be erroneously detected as a novel community.

##### 1.2.2 Shift and return (fig. S19B, H, N)

This matrix represents a sudden shift to a different composition, followed by a period of stability in the second state, then a sudden return to the initial state. We set the abundance of the first six species to zero for the first six and last nine time points to represent the first state. We set the abundance of the seventh to eighteenth species to zero for the seventh to eleventh time points, to represent the second state. We expected to identify the initial shift to the second state as a novel community, but not the subsequent return to the first state.

We detected the shift to the second state as a novel community, and the return as instantaneous novelty. This test highlights the primary advantage of our paired dissimilarity tests. We can reliably identify and separate shifts to unobserved states, and shifts to previously-observed states.

##### 1.2.3 Slow turnover (fig. S19C, I, O)

This matrix represents compositional change from a slow press disturbance, such as warming. Changes in composition are small at each time point but aggregate over time. We accomplished this by converting the lower triangular of our template matrix to zeros. This acts as sequential local extinctions which add up as the time series progresses.

As expected, we found no novel communities in this time series. This test highlights how our expectations capture normal turnover rates, and avoid classifying these as novel communities.

##### 1.2.4 Two-state oscillation (fig. S19D, J, P)

This matrix represents a system that shifts between two alternative states from time point to time point. We set the abundance of the first six species to zero on every odd time point, and set the abundance of the seventh to twelfth species to zero on every even time point. This meant the two states still shared some species (i.e. dissimilarities between states were less than one). We expected to identify the first shift from one state to another as a novel community, but none of the remaining shifts. We detected the first transition as a novel community. Most subsequent transitions were identified as instantaneous novelty, but not cumulative novelty. This is evidence that the dual-nature of our two dissimilarity tests gives us a highly nuanced understanding of community change. We can still identify profound community shifts, separate from those that are shifts to a previously unobserved composition. This test is also evidence that the beta distribution is useful to establish our expectations.

Our prediction intervals are linear on the logit scale, but compressed on a dissimilarity scale as they approach zero and one (e.g., see time points >60 in the bottom panel of fig. S20). In this case, it means we can still detect very large community change as exceeding our expectations, even in a time series built of nothing but large change. This means our expectations are adaptive and track with community dynamics, but have limits, beyond which outliers are still identifiable.

##### 1.2.5 Progressive three-state shift (fig. S19E, K, Q)

This matrix represents two sudden shifts along a gradient, such as two sets of rapid extinctions. In this scenario, state two was more similar to state one than state three. We set the abundance of the seventh to twelfth species to zero for the seventh to twelfth time point (representing state two). We set the abundance of the seventh to eighteenth species to zero for the thirteenth to twentieth time points (representing state three). We expected that both shifts would be detected as shifts to a novel community.

We detected the shift to the second and third state as novel communities. This is evidence that a shift to a novel community does not bias expectations further along the time series, allowing for the detection of multiple, different, novel states in a single time series.

##### 1.2.6 Two shifts and return (fig. S19F, L, R)

In this scenario, the community rapidly shifts to a second state, then immediately to a third, different state, before returning to the initial state. We set the abundance of the first to eighth species to zero for the tenth time point, and the ninth to sixteenth species for the eleventh time point. We expected to identify both shifts as novel communities.

We detected the shift to the second and third state as cumulative dissimilarity, but only the shift from the second to the third as instantaneous dissimilarity (and therefore a novel community). This test highlights that in short time series, having several high dissimilarity values close together in time are sufficient to change our locally-weighted expectations. This was not the case where there were only two high values (in the cumulative dissimilarity test). Time series much longer than 20 time points would need a longer period of high values to adjust expectations, and in that case, as we have defined ‘novelty’ relative to expectations from the local time window, we would correctly exclude these high values as they emerged during a prolonged period of high dissimilarity. This is evident in the two-state oscillation section of the combination test below.

##### 1.2.7 Combination (fig. S20)

Given that real time series likely contain a variety of compositional change over time, we combined the above scenarios into a single, 120 point time series to see how our framework operated on a longer time series with various dynamics.

When we combined various community dynamics together, we observe many of the same patterns as when the time series were separated. Most of the same shifts were identified as novel communities, except where we re-used the same ‘state’ in multiple tests (such as the second shift in the ‘progressive three-state shift' section towards the end. This combined time series shows an example of how the locally-weighted expectations change along the time series. The expectations for instantaneous dissimilarity rose during the period of a two-state oscillation. This affected our expectations for the first shift in the 'progressive three-state shift'. The expectations for cumulative dissimilarity remained stable during the the beginning of the 'slow turnover' because of the lower values in the 'two-state oscillation' section.

#### 1.3 Testing our framework using a random walk simulation

We simulated random walk time series, parameterized with the demographic models used to create Fig. 5 in the main text. (see Methods for model details). The simulation was designed as a 'null' model, to identify how often we expected novel communities to arise if taxa within simulated communities were allowed to turnover at probabilities observed in the NSB dataset. This was designed to test whether the results we obtained in the observed NSB data were similar to those we would be expect from random-walk communities. The simulation process used the following steps.

1. We initialized 9,999 time series, drawing their length (number of time bins) and gamma diversity (total number of species observed along the time series) from Poisson distributions generated from the observed NSB data.

2. For each time series, we seeded the time series with an initial set of species, drawing the alpha diversity at the first time point from a Poisson distribution generated from the alpha diversities observed in the NSB data.

3. We then allowed species presence and absence to iterate at each time point, using two rules:

   (a) We classified each time point as 'instantaneous novelty', 'cumulative novelty' or a 'novel community', with the probability of each drawn from binomial distributions populated from the 'random-taxa' models used to generate Fig. 1C. If none of these classifications were drawn from the binomial distributions, the time point was classified as 'background'. If multiple classifications were drawn (as they were drawn from independent probability distributions), we repeated the binomial draw until there was only one classification for the time point. Within each time point:

      - present species could emigrate
      - non-originated species could originate
      - present, originated species could go locally extinct
      - absent, originated and non-extinct species could re-immigrate.

   (b) The probability of each of these events occurring for each relevant taxa was drawn from binomial distributions generated using probabilities from demographic models, using the probabilities for the time point's classification. See Methods for details on the demographic models.

4. We ran each simulated time series through our novel community detection framework.

5. We modeled the probability of each of our four community classifications occurring across our simulated time series, treating each simulation as a set of weighted trials, using the same model syntax as the models used to estimate expected classification probability in Fig. 1C.

We found that the probability of identifying cumulative novelty and novel communities were more likely in the null model than in the NSB data, and instantaneous novelty was less likely (table S12). This suggests that in real ecological time series, the emergence of new compositions tends to occur less often than expected by chance. We believe this result is due to how our simulation is set up. In our simulation, the probability of taxonomic turnover is calculated separately for each taxon, which means that any synchronized extinction or origination across taxa is only due to random chance. In real time series, we may be seeing a greater amount of synchronization of taxonomic turnover, whereby multiple taxa originated or went extinct in a single bin. This could be due to shared environmental or competitive tolerances, for instance. Synchronized turnover events would have reduced the number of novel compositions observed relative to if they each went extinct in a separate bin. We noted initially that the probability of instantaneous novelty was lower in the null simulation (table S12), but novel communities also had to be classified as instantaneous novelty. This meant that there was a combined 5.13% chance of instantaneous novelty or novel community in the simulation, and 5.73% in the observed data, which, given the uncertainty, is similar.

#### 2 Supplementary figures

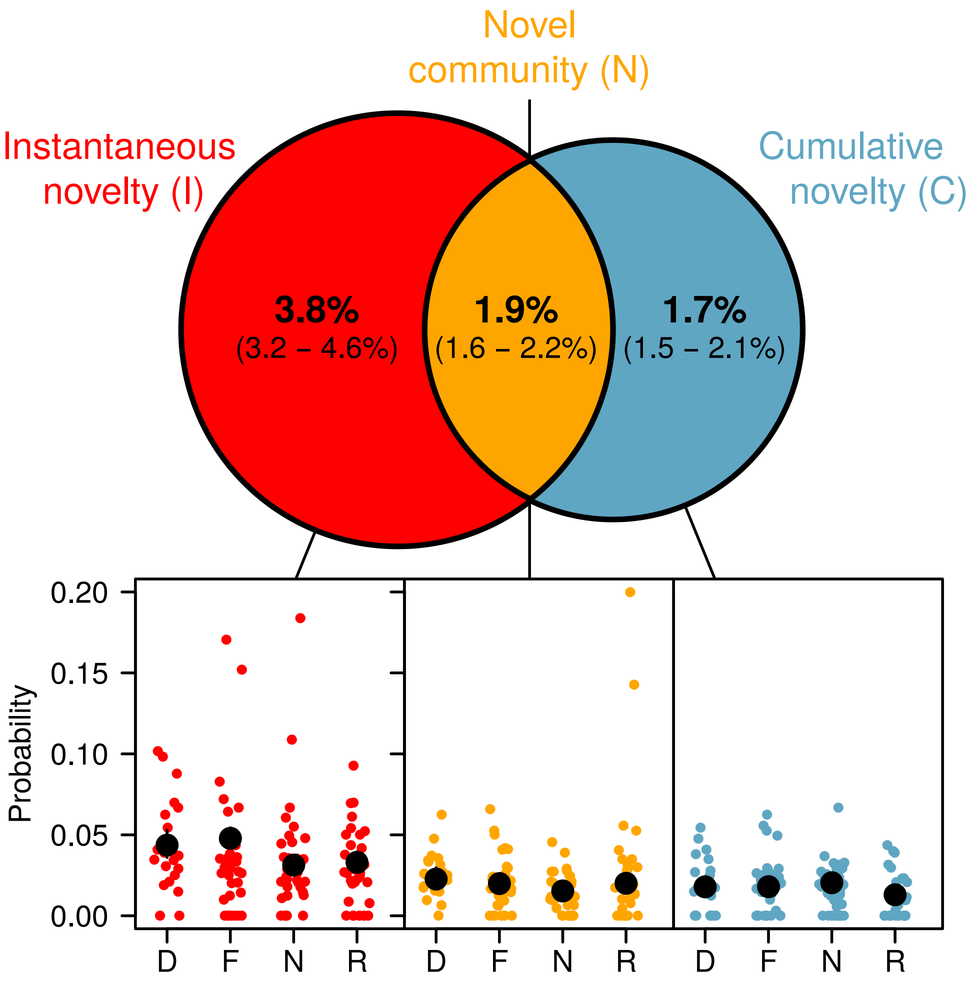

**Figure S1.** Probability of community classifications using our novel community detection framework. This version identified novel communities using individual core time series, rather than aggregated at the Longhurst biogeographical province scale. The Venn diagram shows the estimated probability of a community being classified as either 'instantaneous novelty' (exhibiting only a large shift in composition), or 'cumulative novelty' (exhibiting only a shift to a new state), or as a 'novel community' (exhibiting both). Probabilities were derived from logistic regressions. Probability ranges indicated on the Venn diagrams are regressions treating the four planktonic groups as levels of a random effect (percentages in brackets are 95% confidence intervals (CI)). Each set of percentages represents a single model. Sub-plots are also separate models, treating planktonic taxa groups as a fixed categorical variable (errors bars are 95% CIs). D = diatoms, F = foraminifers, N = calcareous nannoplankton and R = radiolarians. Colored points in subplots represent the proportion of each Longhurst Province, for each taxon group, classified as novelty.

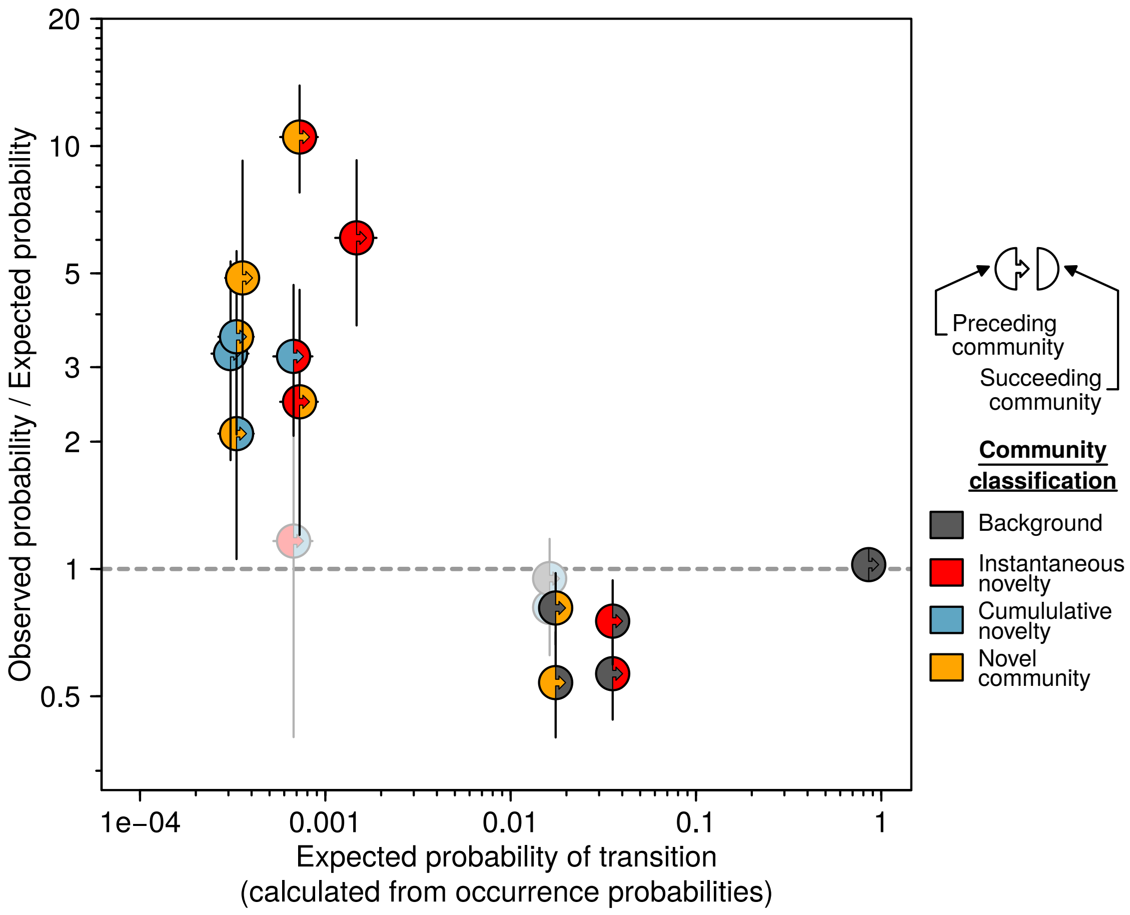

**Figure S2.** Expected probability of transitions between each of our four community classifications, and the ratio of observed transition probabilities to expected probabilities. This version identified novel communities using individual core time series, rather than aggregated at the Longhurst biogeographical province scale. Expected transition probabilities were estimated by multiplying the probabilities of occurrence for the preceding and succeeding classification from occurrence models. Observed transition probabilities were estimated using binomial generalized linear mixed-effect models. Points are halved and dual-colored; the left-hand and right-hand color represent the classification of the preceding and succeeding community, respectively. Y-axis points are ratios: values greater than one indicate transition probability was higher in observed data than expected, and vice versa. Colored points are transitions where observed to expected ratio was significantly different to one: faded points had confidence intervals that crossed one. Note that both axes are ln-transformed.

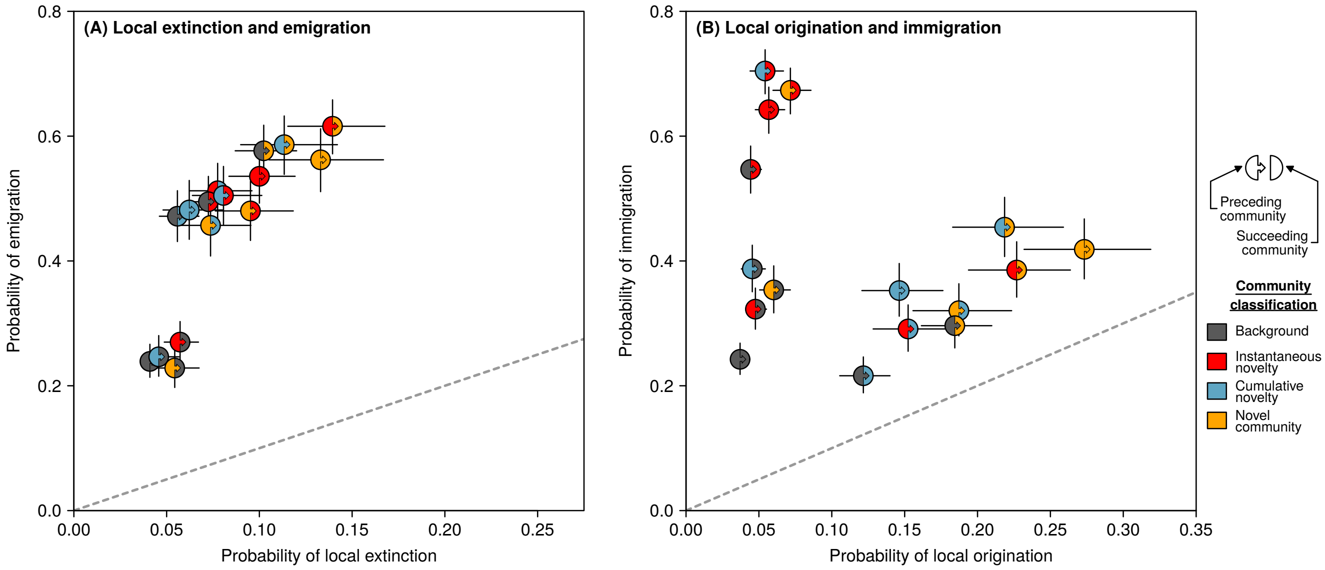

**Figure S3.** Probability of planktonic taxa exhibiting one of four demographic changes in the transition between two communities along a time series. This version identified novel communities using individual core time series, rather than aggregated at the Longhurst biogeographical province scale. Taxonomic gain plotted as mean predictions from generalized linear mixed-effects models, one for each demographic change, fit with random intercepts for time series nested within planktonic taxa groups. **(A)** Taxonomic loss plotted as the probability of local extinction (taxon disappears from time series and does not return) against the probability of emigration (species disappears transiently). **(B)** probability of local origination (taxon appears in time series for the first time) against the probability of immigration (subsequent reemergence of a previously-present taxon). Points are halved and dual-colored; the left-hand and right-hand color represents the classification of the preceding and succeeding community, respectively. Predictions were obtained from four generalized linear mixed-effects models The dashed line shows a 1:1 ratio. Model summaries are found in tables S6-S11.

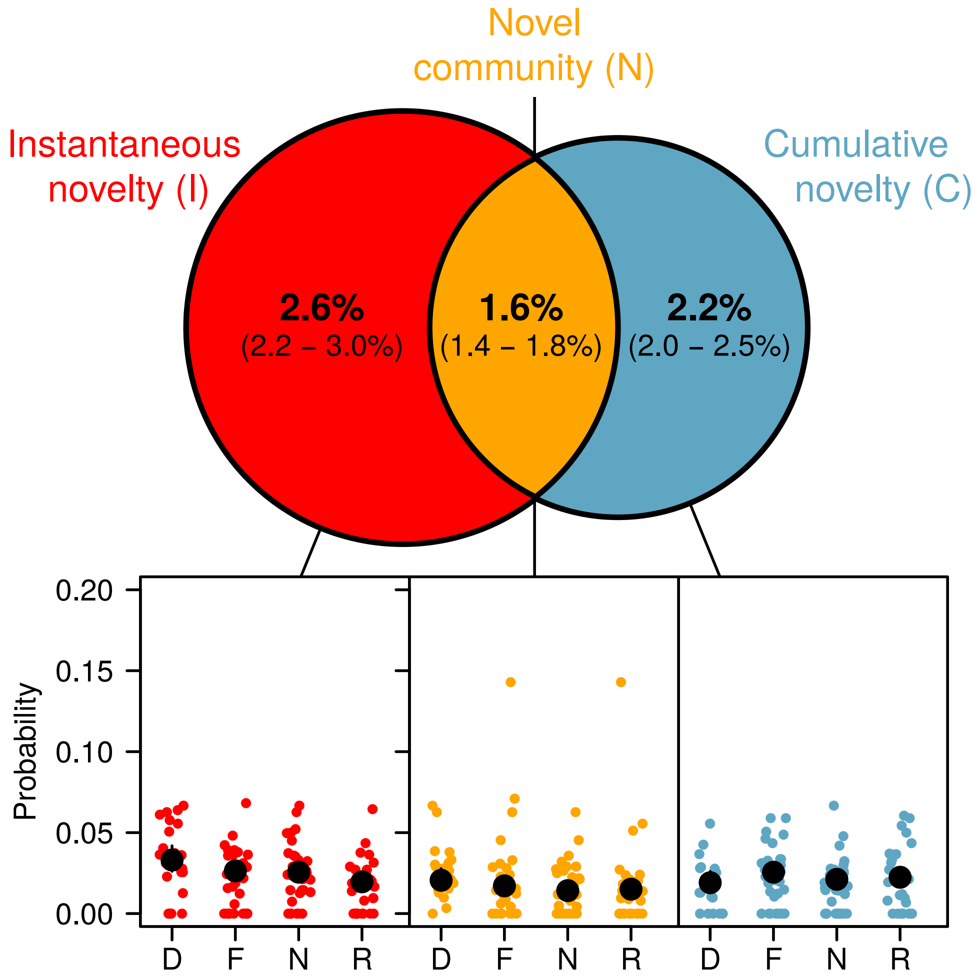

**Figure S4.** Probability of community classifications using our novel community detection framework. This version was conducted with data aggregated at the Longhurst scale (as per the main text figures), but excludes time series sampling bins with fewer than five species. See fig. S1 caption for other figure details.

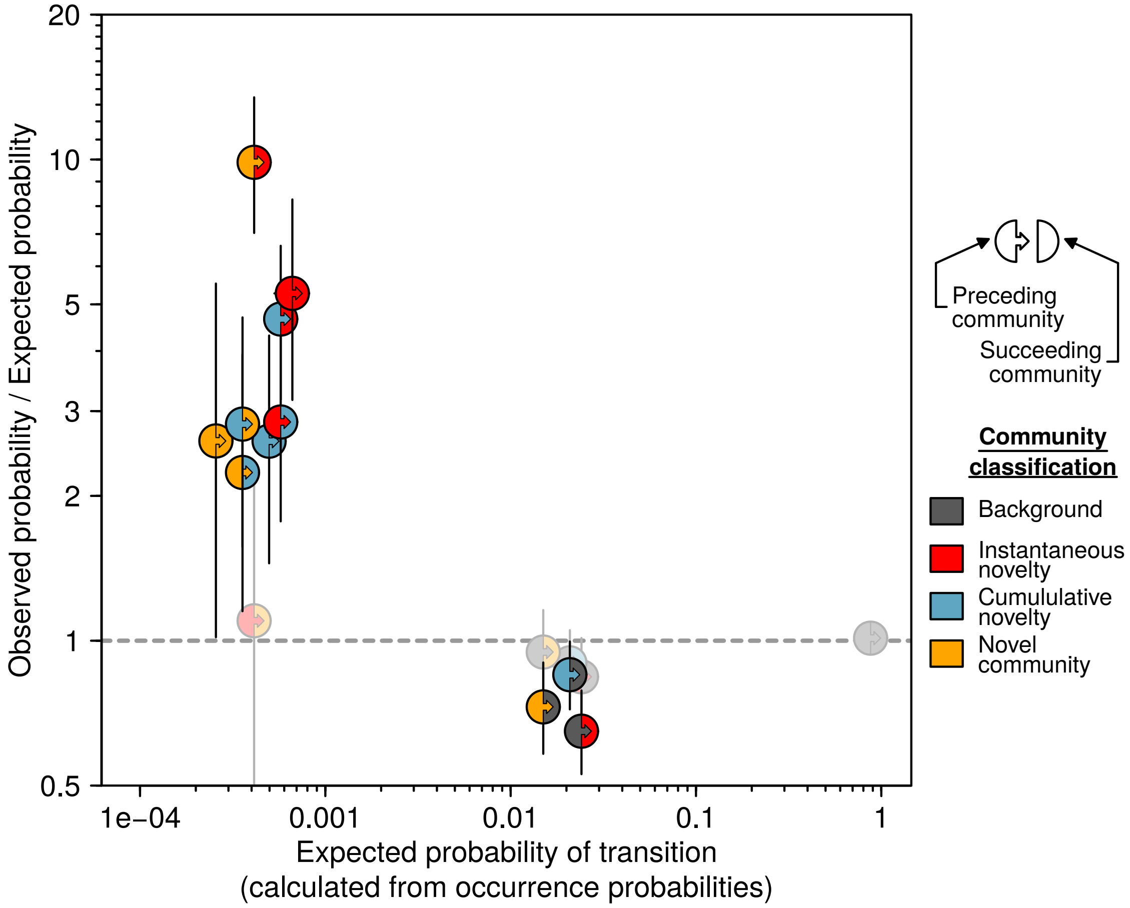

**Figure S5.** Expected probability of transitions between each of our four community classifications, and the ratio of observed transition probabilities to expected probabilities. This version was conducted with data aggregated at the Longhurst scale (as per the main text figures), but excludes time series sampling bins with fewer than five species. See fig. S2 caption for other figure details.

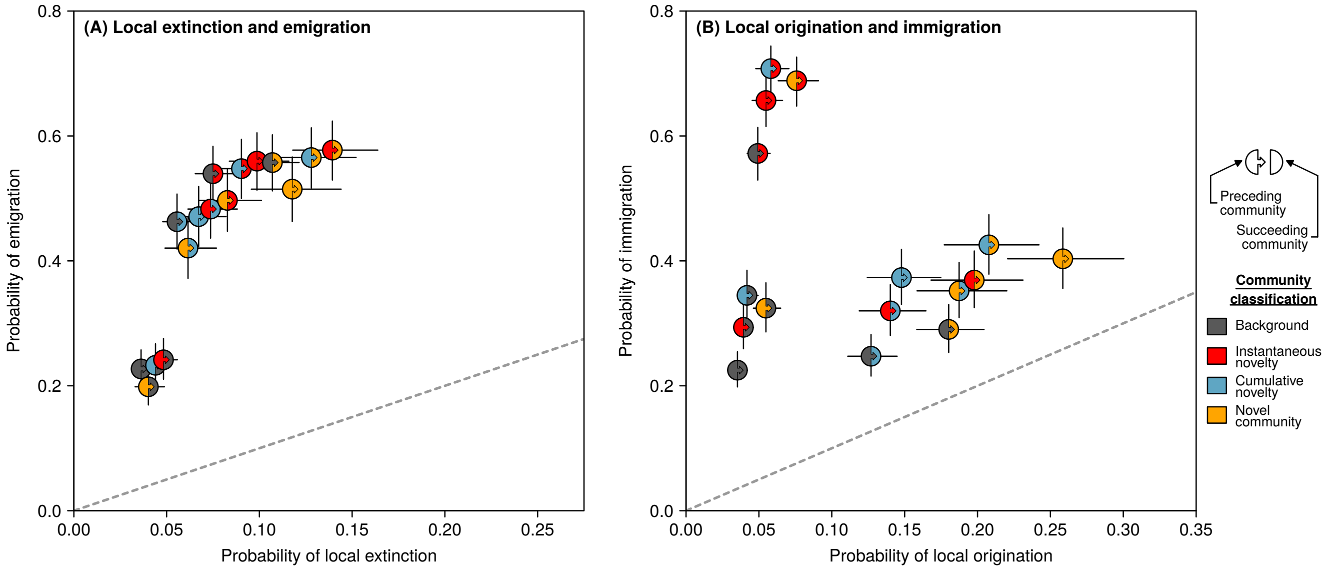

**Figure S6.** Probability of planktonic taxa exhibiting one of four demographic changes in the transition between two communities along a time series. This version was conducted with data aggregated at the Longhurst scale (as per the main text figures), but excludes time series sampling bins with fewer than five species. See fig. S3 caption for other figure details.

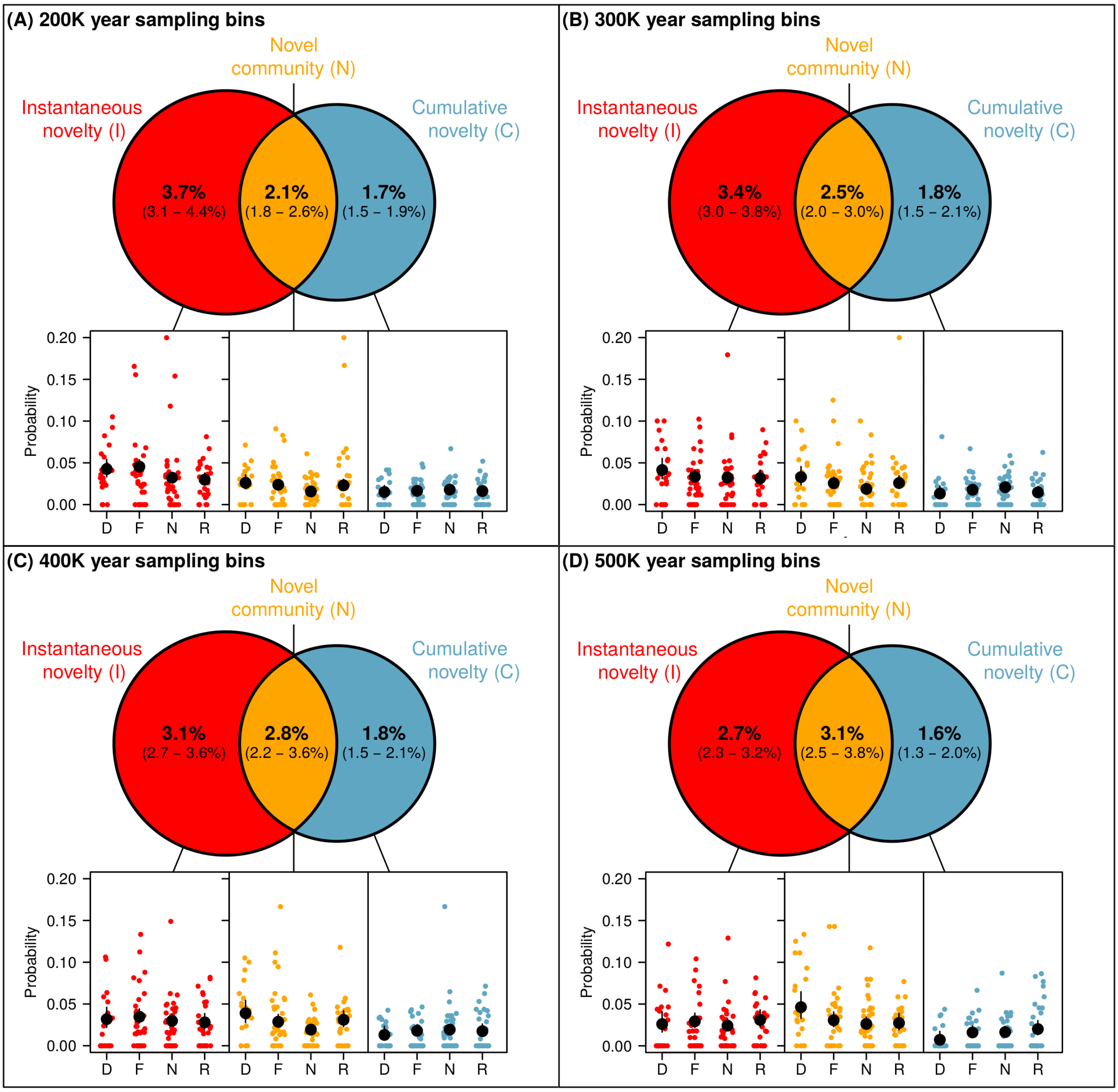

**Figure S7.** Probability of community classifications using our novel community detection framework. This version was conducted with data aggregated at the Longhurst scale (as per the main text figures), but uses different time windows to aggregate NSB data for novel community detection. See fig. S1 caption for other figure details.

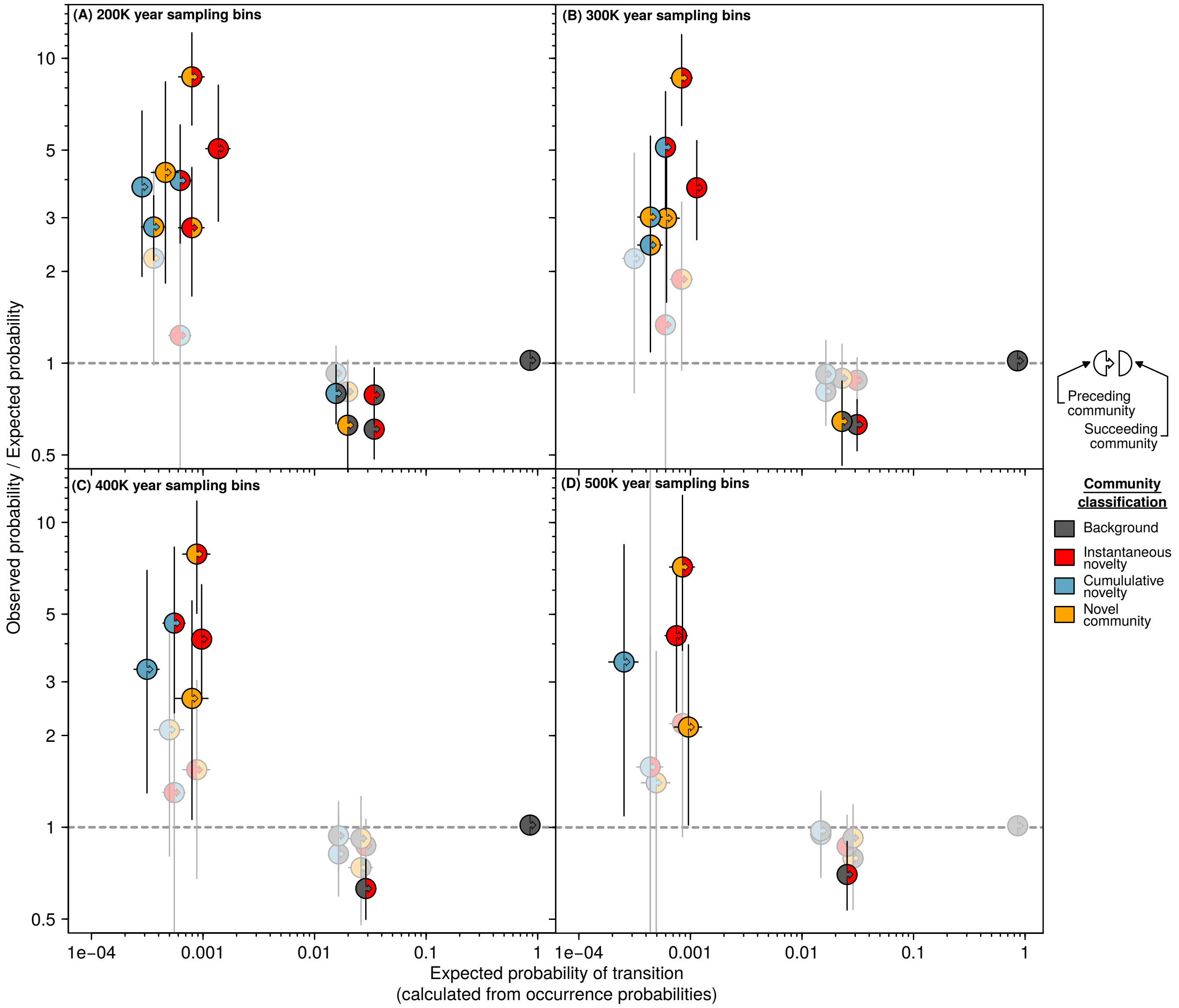

**Figure S8.** Expected probability of transitions between each of our four community classifications, and the ratio of observed transition probabilities to expected probabilities. This version was conducted with data aggregated at the Longhurst scale (as per the main text figures), but uses different time windows to aggregate NSB data for novel community detection. See fig. S2 caption for other figure details.

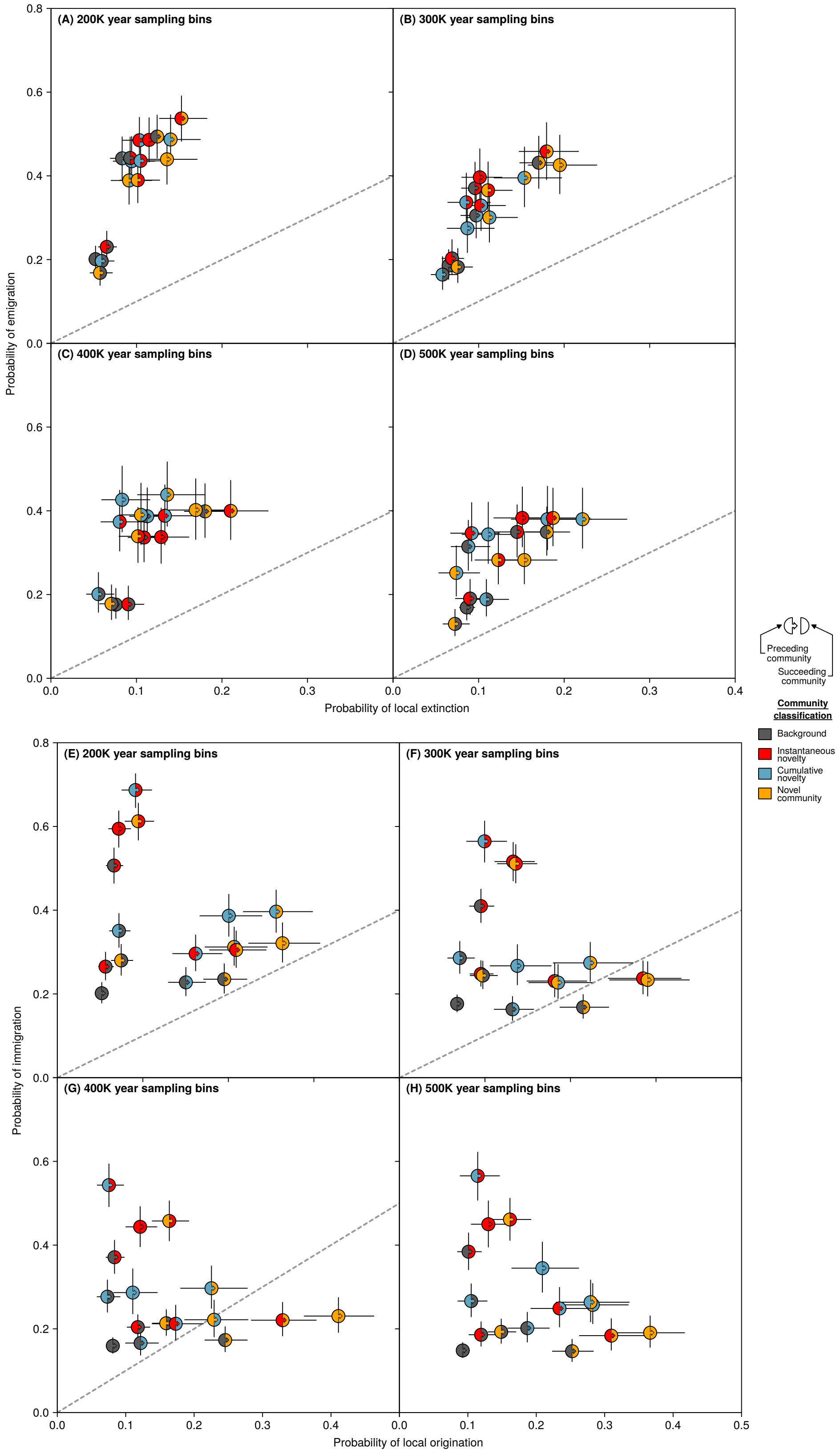

**Figure S9.** Probability of planktonic taxa exhibiting one of four demographic changes in the transition between two communities along a time series. This version was conducted with data aggregated at the Longhurst scale (as per the main text figures), but uses different time windows to aggregate NSB data for novel community detection. See fig. S3 caption for other figure details.

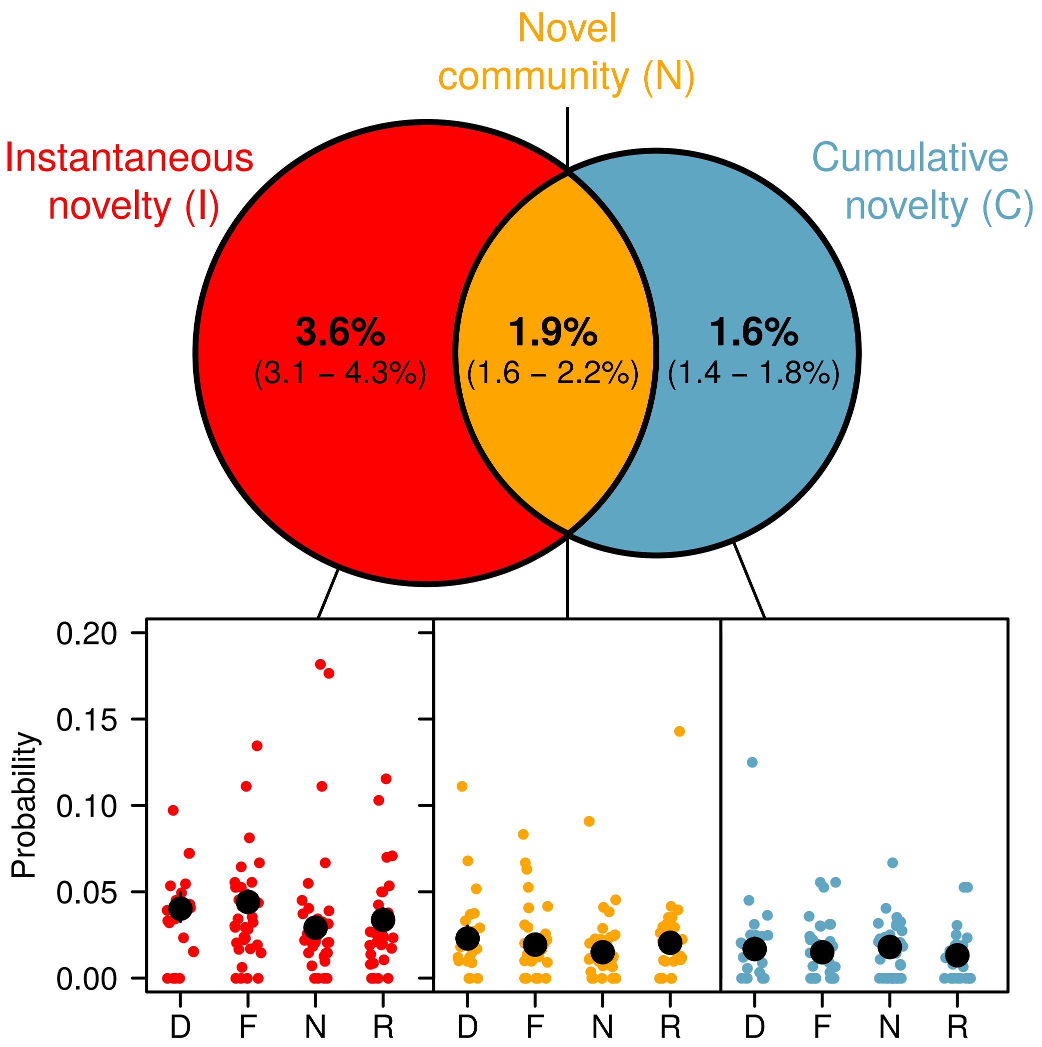

**Figure S10.** Probability of community classifications using our novel community detection framework. This version was conducted with data aggregated at the Longhurst scale (as per the main text figures), but removes the bottom and top 5% of each taxon's occurrences in the Neptune Sandbox ("Pacman" analyses). See fig. S1 caption for other figure details.

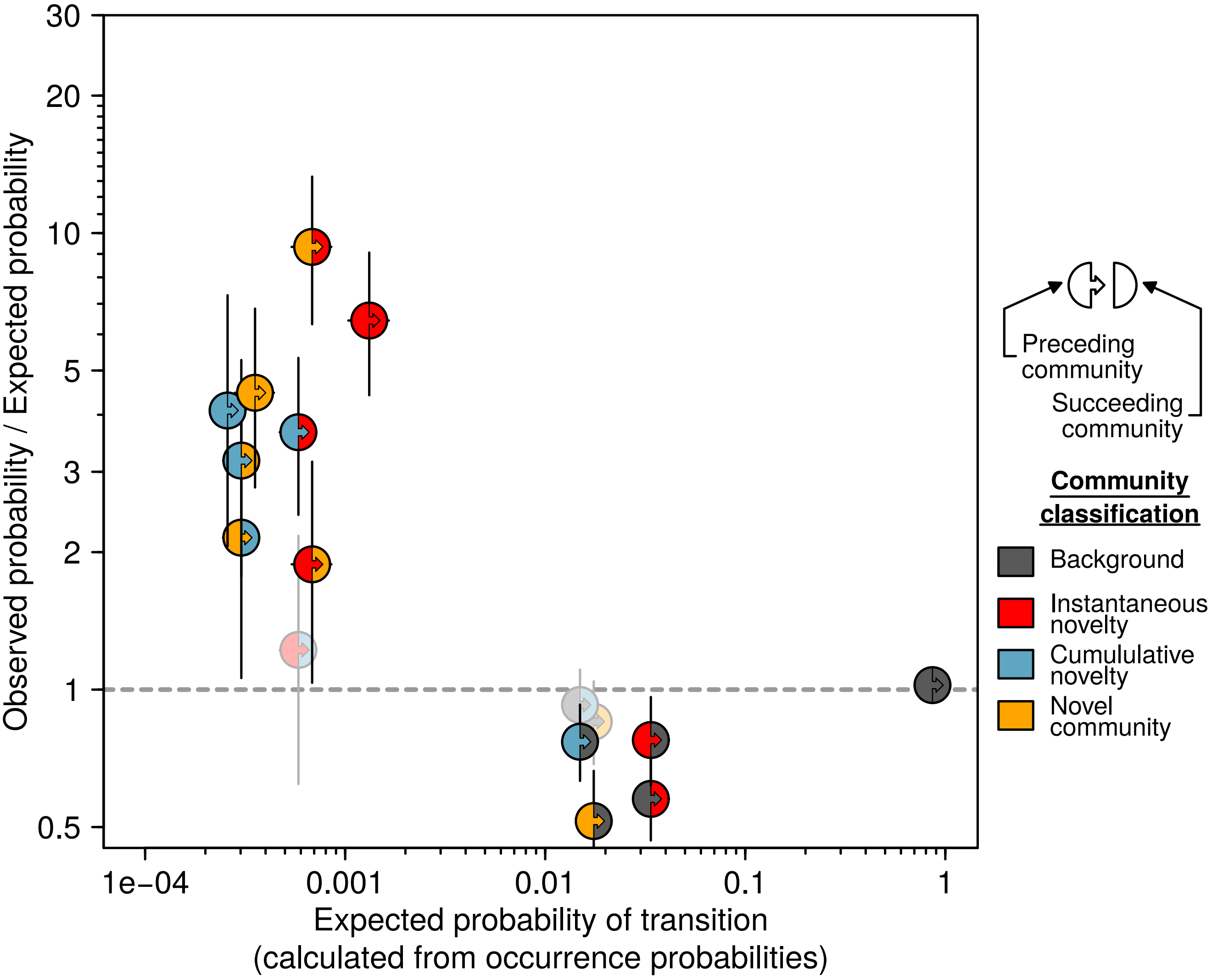

**Figure S11.** Expected probability of transitions between each of our four community classifications, and the ratio of observed transition probabilities to expected probabilities. This version was conducted with data aggregated at the Longhurst scale (as per the main text figures), but removes the bottom and top 5% of each taxon's occurrences in the Neptune Sandbox ("Pacman" analyses). See fig. S2 caption for other figure details.

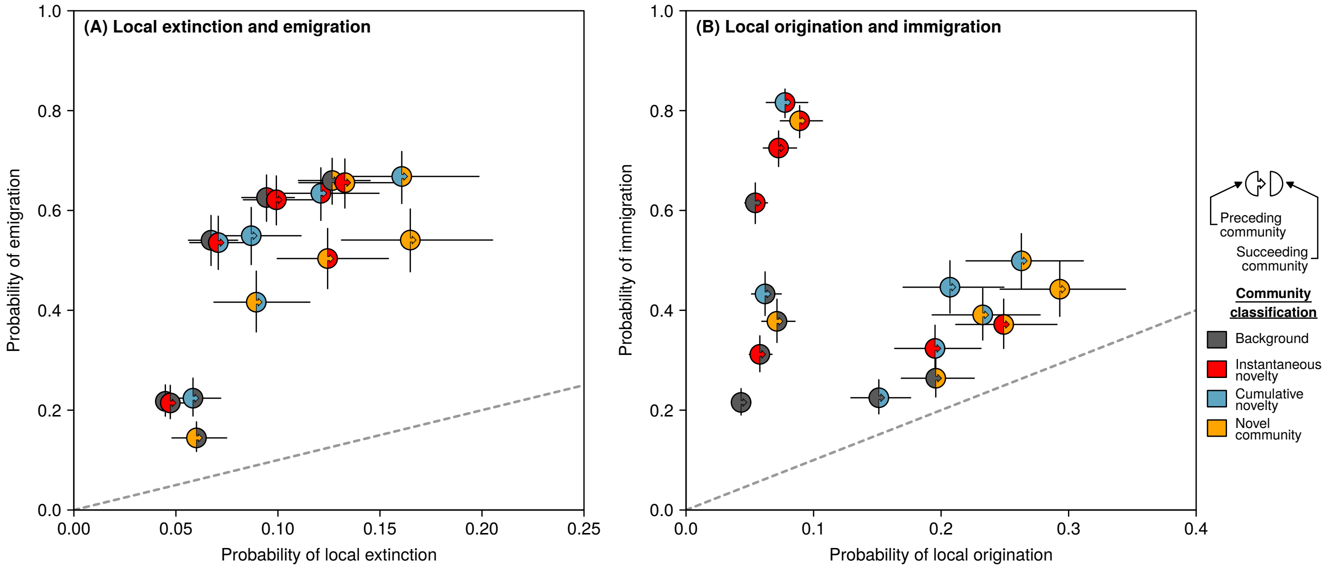

**Figure S12.** Probability of planktonic taxa exhibiting one of four demographic changes in the transition between two communities along a time series. This version was conducted with data aggregated at the Longhurst scale (as per the main text figures), but removes the bottom and top 5% of each taxon's occurrences in the Neptune Sandbox ("Pacman" analyses). See fig. S3 caption for other figure details.

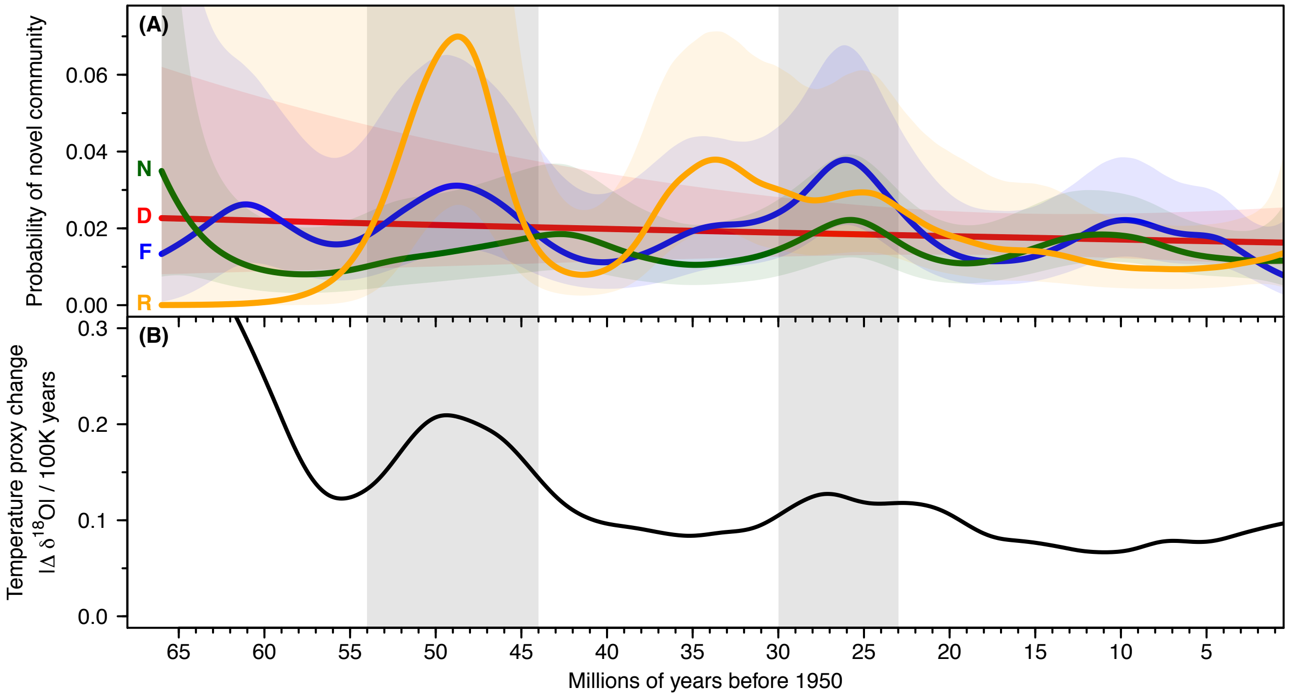

**Figure S13.** Temporal trends in the **(A)** emergence probability of novel communities, separately for calcareous nannoplankton ("N", green), diatoms ("D", red), foraminifera ("F", blue) and radiolarians ("R", yellow), and **(B)** trend in temperature change over time, using the $\delta^{18}O$ ratios from (*43*). Shaded regions are 95% confidence intervals of mean trends. Grey rectangles are two regions where novel community emergence peaked alongside increases in temperature change.

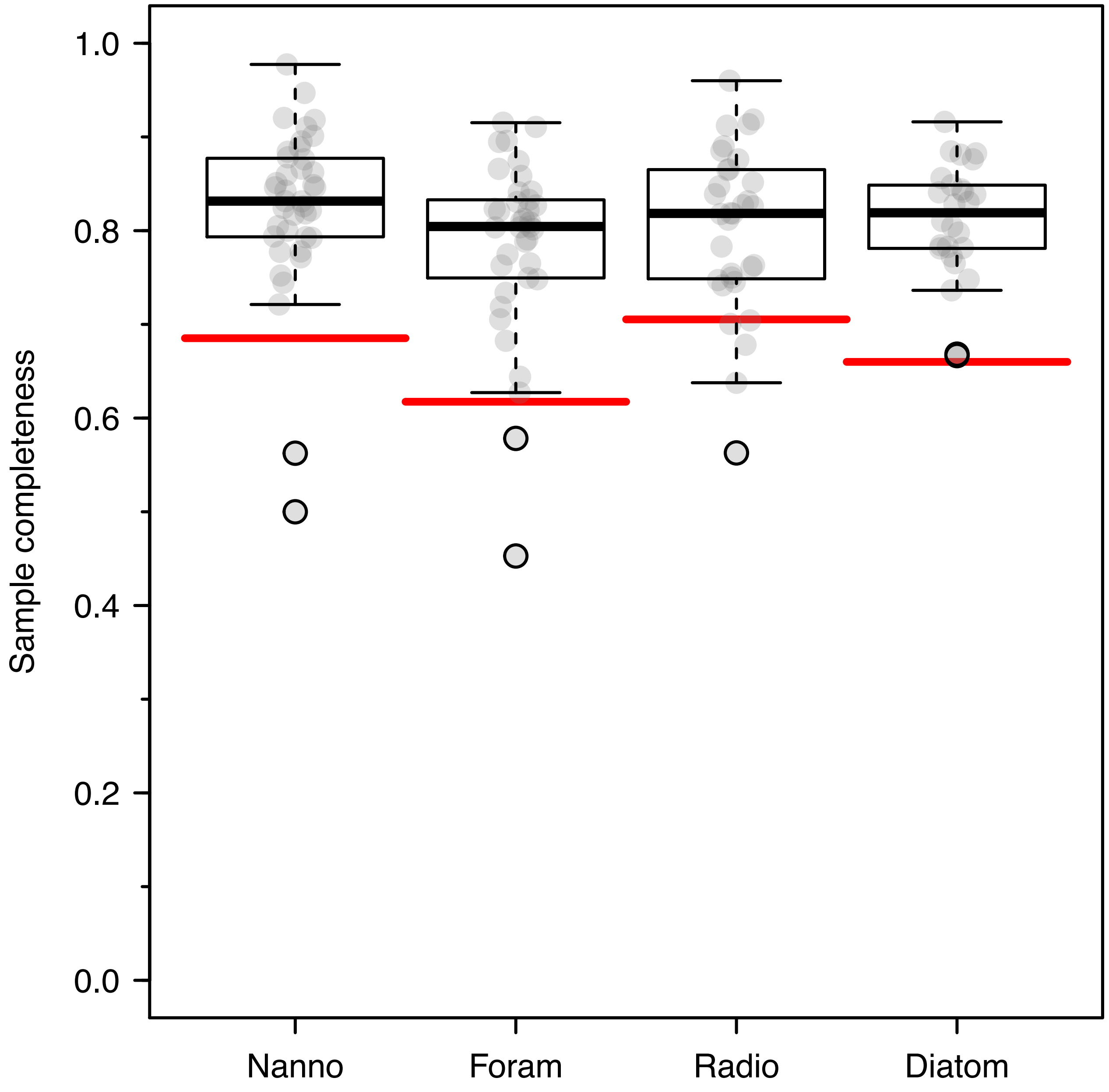

**Figure S14.** Sample completeness of Longhurst province time series, as assessed by the ‘three-timer’ methodology (*27*). This method uses locally-weighted sampling probability to assess sample completeness among samples. The red horizontal lines are the lower 5% quantile. Time series below this point were considered to be potentially taphonomically-biased and were excluded from subsequent analyses.

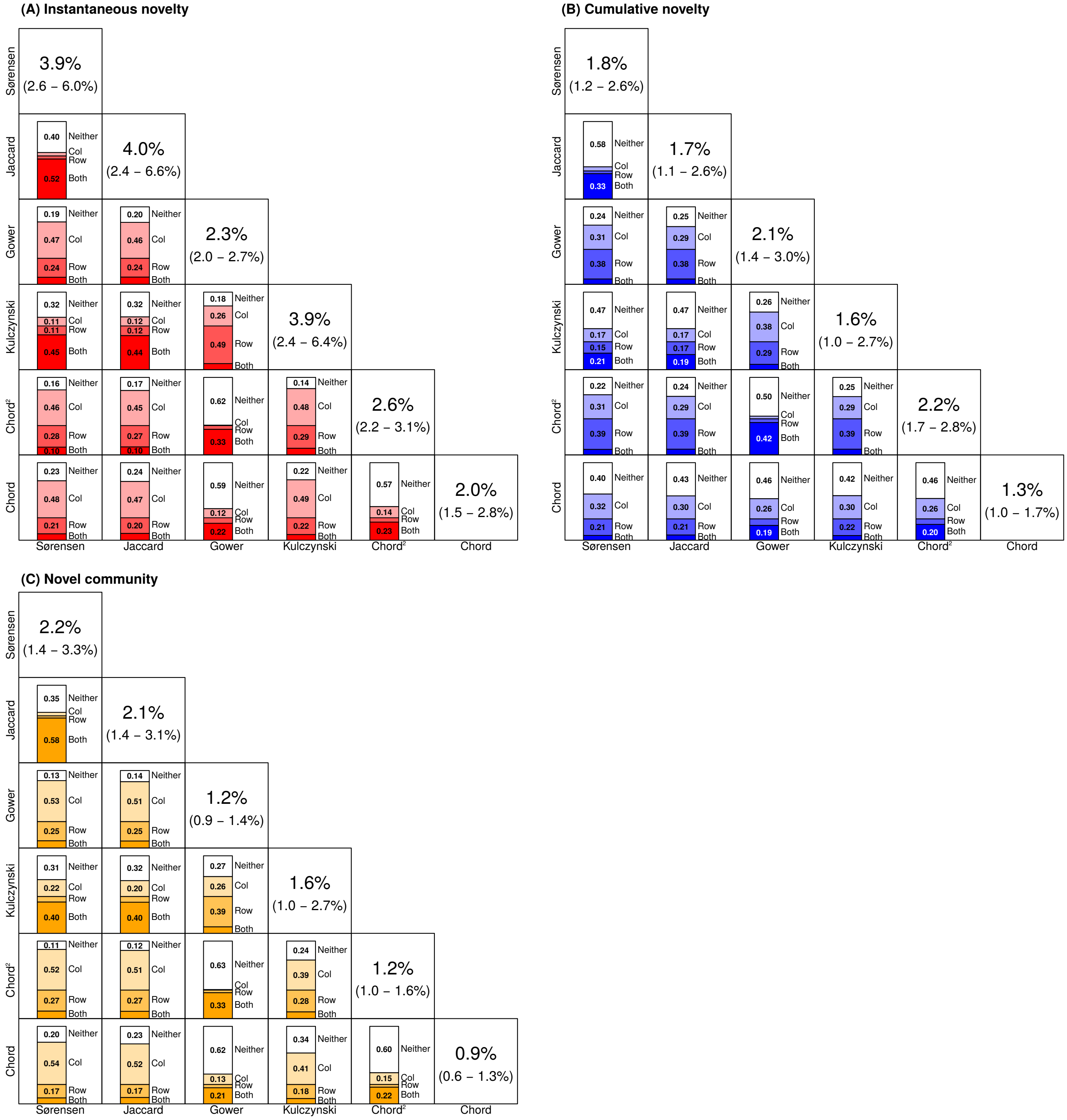

**Figure S15.** Comparison of our novel detection framework classifications using six different dissimilarity indices for the calcareous nannoplankton dataset. **(A)** Instantaneous novelty, **(B)** cumulative novelty, and **(C)** novel communities. Panel diagonals show the proportion of communities identified as each community classification. Off-diagonals show the number of communities classified using both dissimilarity indices ("Both"), just the row dissimilarity index ("Row"), just the column dissimilarity index ("Col"), or identified by neither ("Neither"). The proportions sum to one, and represent all communities identified by any of the six dissimilarity indices. As an example, in (C), Jaccard vs Sørensen has a "Neither" proportion of 0.35. These two indices were very similar ("Row" and "Col" are very small), and the "Neither" represents novel communities identified by the other indices that were not identified by either the row (Jaccard) or column (Sørensen) index.

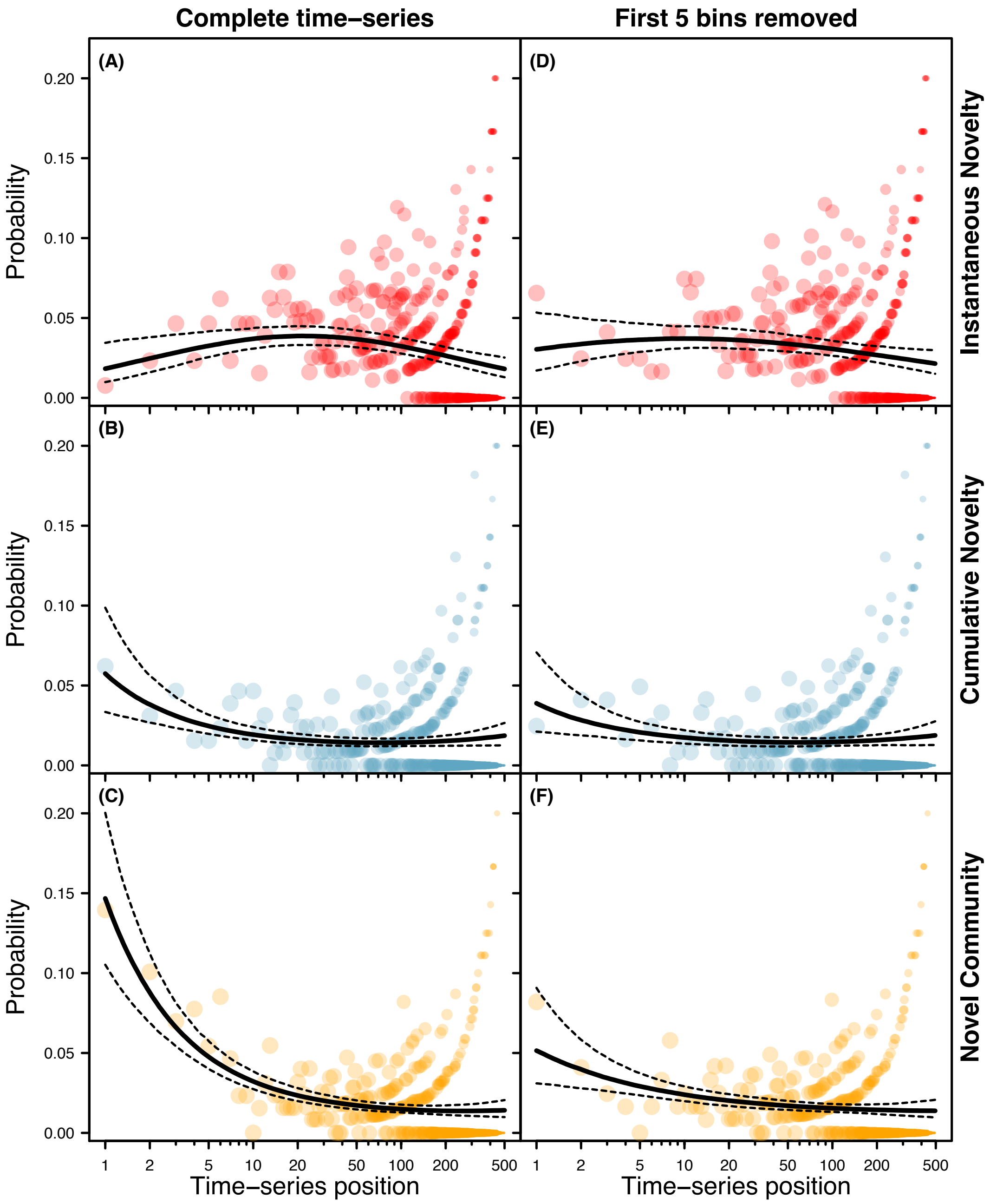

**Figure S16.** The probability of detecting novelty depending on position within time series. Points are the proportion of bins at each position classified, with the size of the point proportional to the number of time series that have a bin at that position (sample sizes reduce as time series position increases as there were few time series longer than 150 bins). Lines are modeled probabilities and 95% confidence intervals. **(A-C)** shows probabilities when the entirety of each time series was included; **(D-F)** shows probabilities after the first five bins of each time series were excluded. Positions were relabelled for these models: position one in A-C is the 2nd bin in the time series (i.e the first bin we could calculate instantaneous and cumulative dissimilarity values for). Position one in D-F is equivalent to position five in A-C, and position six in the raw time series data.

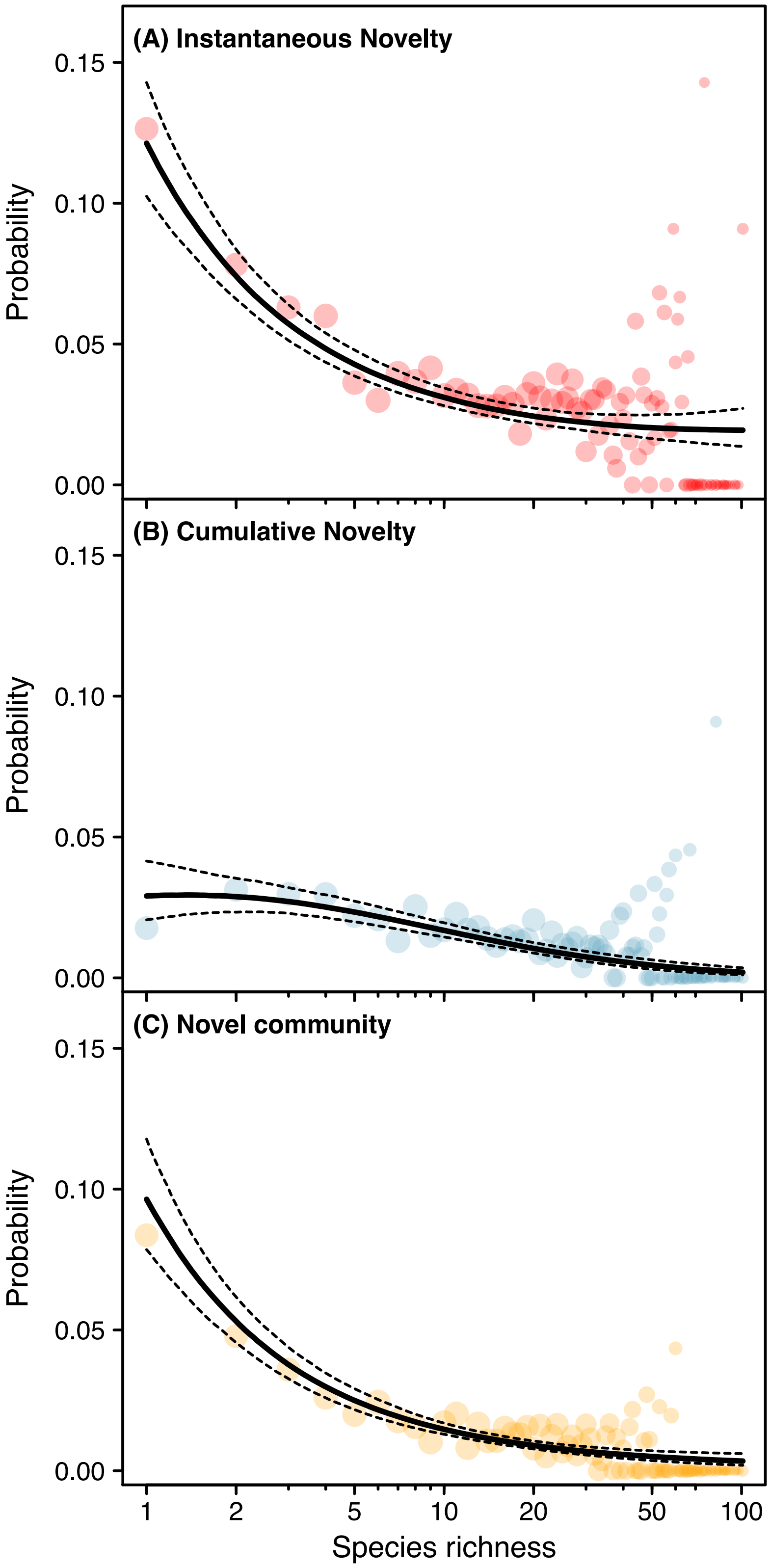

**Figure S17.** The probability of classifying a community as **(A)** instantaneous novelty, **(B)** cumulative novelty, or **(C)** a novel community, depending on the species richness of that community. Points are the proportion of bins at each position classified as each community type, with the size of the point proportional to the number of communities with that richness. Lines are modeled probabilities and 95% confidence intervals from binomial generalized linear mixed-effects models.

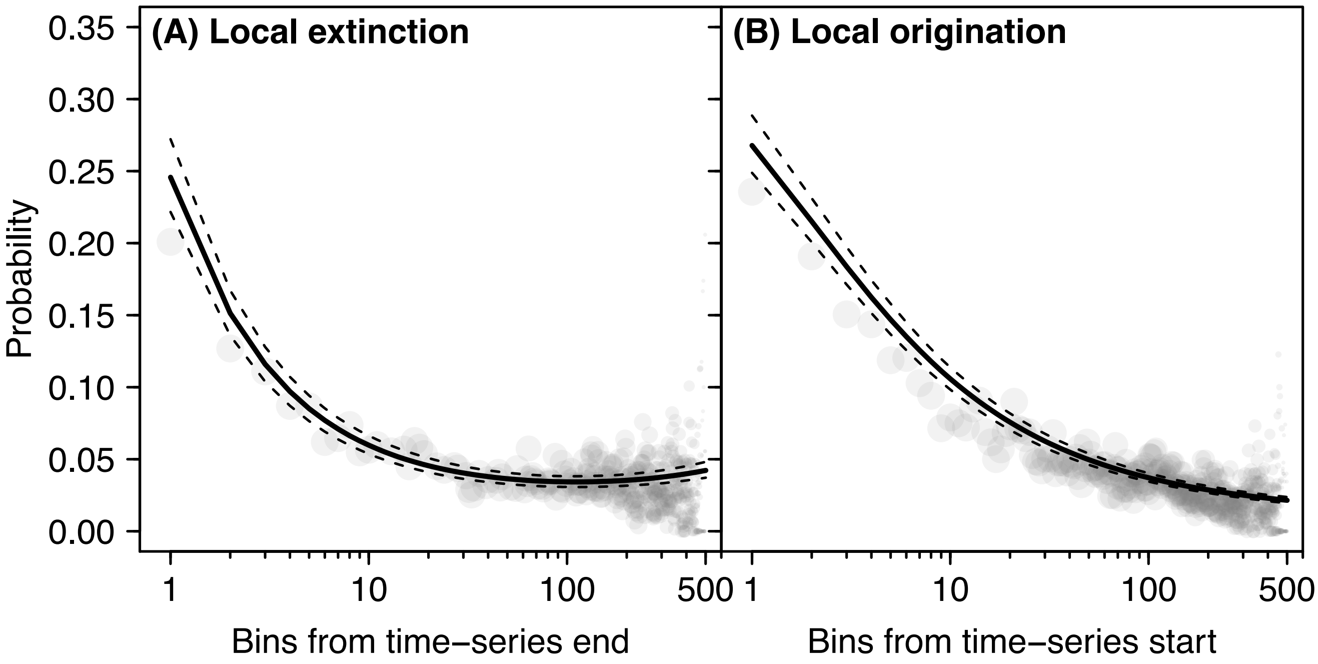

**Figure S18.** Probability of a taxon **(A)** becoming locally extinct, or **(B)** locally originating based on the number of bins from the start or end of the time series, respectively. Lines are modeled probabilities, dashed lines are 95% confidence intervals. Non-linear trends terminate at position ten: we removed these bins from each time-series prior to running our local extinction and origination models. Remaining linear trends were accounted for by including "number of time points from time series start/end" as a model covariate.

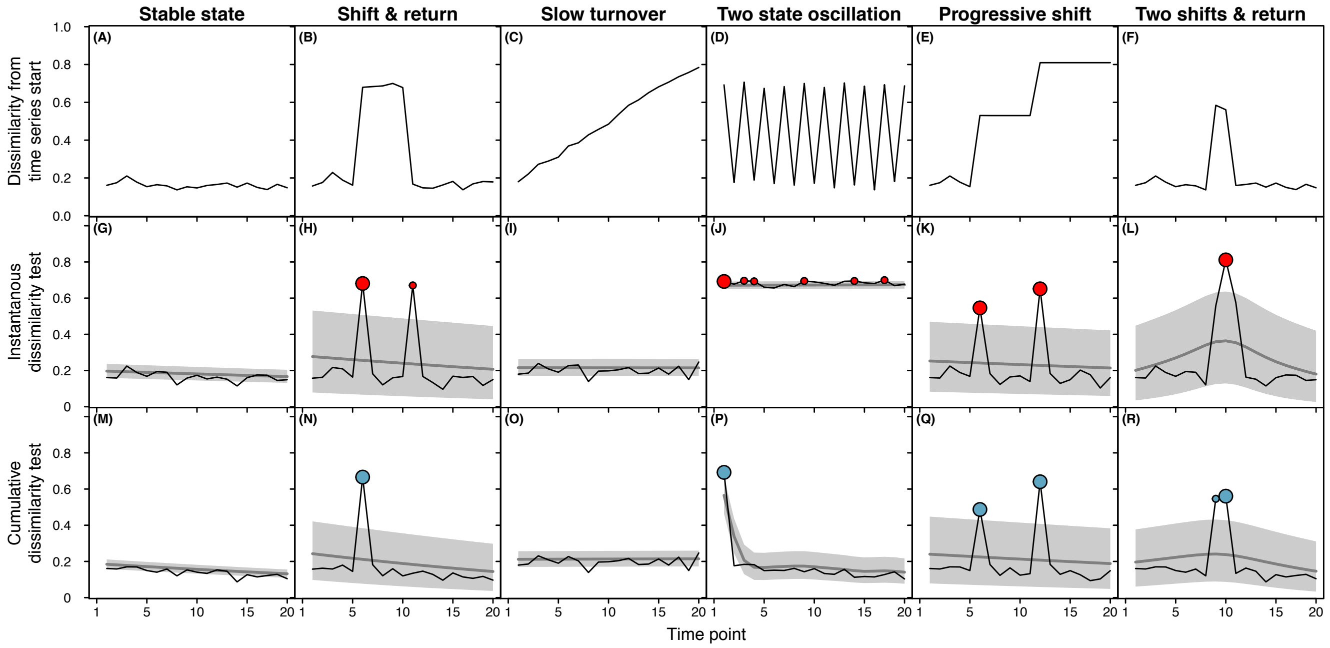

**Figure S19.** Simulations for six different community turnover scenarios, which we used to benchmark the performance of our novelty detection framework. Time series run from time point one (oldest) to time point 20 (youngest), but because they show dissimilarities to past time points, the first value is for time point two. The top row of sub-plots show the dissimilarity along the time series relative to the final time point. The central and bottom row are our two novelty tests. In these sub-plots, the grey line represents mean expected dissimilarity, with the grey polygon showing 95% prediction intervals. Time points above this threshold were classified as instantaneous or cumulative novelty. Larger points are those classified as both instantaneous and cumulative novelty, which qualify as a 'novel community' under our framework. Descriptions of how these time series were generated can be found above in the Supplementary Methods.

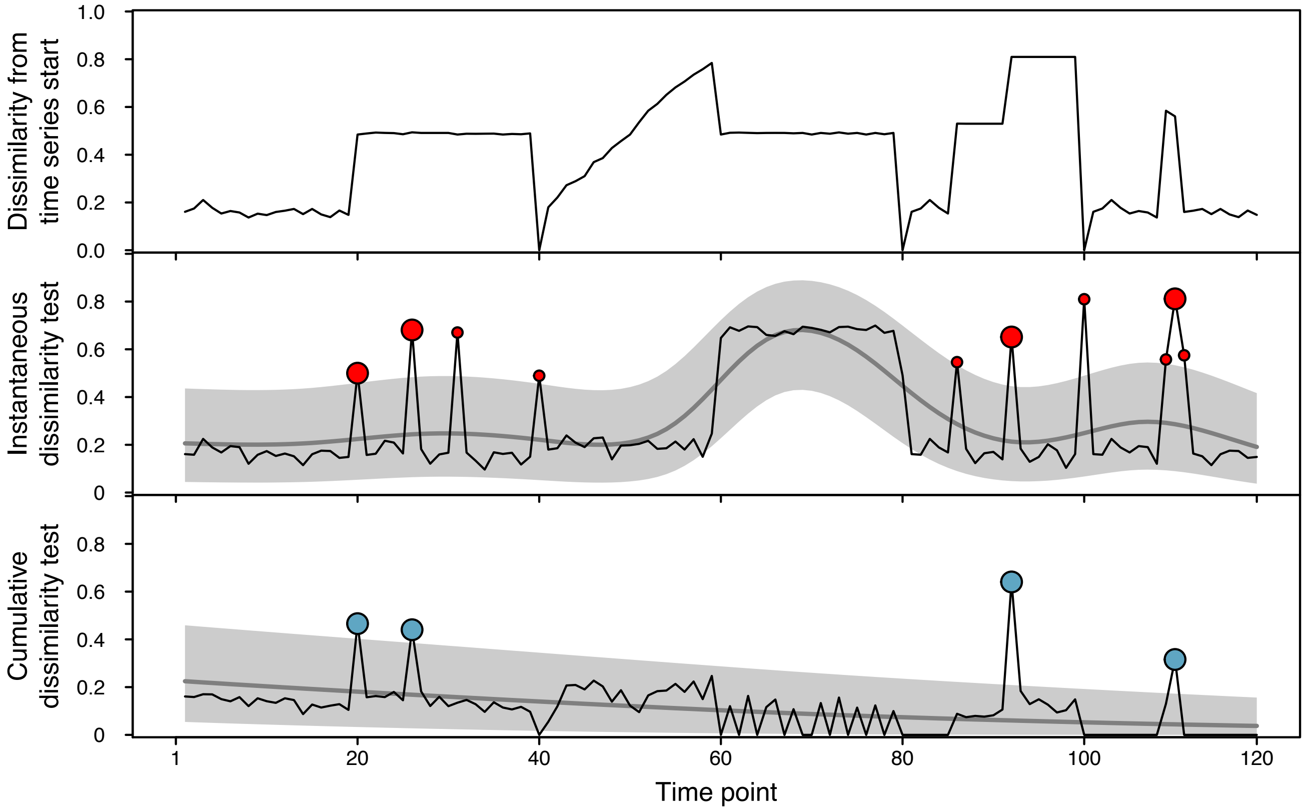

**Figure S20.** A combination of the individual simulated time series applied in fig. S19, benchmarked against our novelty detection framework. Time series runs from point one (oldest) to 120 (youngest), but because they show dissimilarities to past time points, the first value is for time point two. The top sub-plot shows the dissimilarity along the time series relative to the final time point. The central and bottom subplots are our two novelty tests. In these sub-plots, the grey line represents mean expected dissimilarity, with the grey polygon showing 95% prediction intervals. Time points above this threshold were classified as instantaneous or cumulative novelty. Larger points are those classified as both instantaneous and cumulative novelty, which qualify as a 'novel community' under our framework.

### 3 Supplementary tables

#### 3.1 Occurrence probability model tables

Tables S1-S4 are summaries of models estimating the probability of novelty in the Neptune Sandbox dataset. Table S1 is a summary of three binomial generalized linear mixed-effects models (logit link function), each treating the four planktonic groups as levels of a random intercept (”random taxa models”). Tables S2-4 are generalized linear models (logit link function) treating the four planktonic groups as a categorical fixed effect (“fixed taxa models”). These are interceptless models that generate from-zero slope estimates for each taxa group.

**Table S1.** Summary table of random taxa novelty probability models.

| | Estimate | SE | Z-value | P-value | Random variance |
|---|---|---|---|---|---|
| Novel community | -3.952 | 0.077 | -51.235 | $< 0.001$ | 0.009 |
| Cumulative novelty | -4.029 | 0.084 | -47.827 | $< 0.001$ | 0.012 |
| Instantaneous novelty | -3.225 | 0.096 | -33.516 | $< 0.001$ | 0.030 |

**Table S2.** Summary table of fixed taxa instantaneous novelty probability model.

| | Estimate | SE | Z-value | P-value |
|---|---|---|---|---|
| Diatoms | -3.087 | 0.102 | -30.193 | $< 0.001$ |
| Foraminifera | -2.991 | 0.069 | -43.384 | $< 0.001$ |
| Nannoplankton | -3.432 | 0.071 | -48.247 | $< 0.001$ |
| Radiolarians | -3.377 | 0.096 | -35.148 | $< 0.001$ |

**Table S3.** Summary table of fixed taxa cumulative novelty probability model.

| | Estimate | SE | Z-value | P-value |
|---|---|---|---|---|
| Diatoms | -4.006 | 0.158 | -25.418 | $< 0.001$ |
| Foraminfera | -4.001 | 0.111 | -36.122 | $< 0.001$ |
| Nannoplankton | -3.871 | 0.088 | -44.185 | $< 0.001$ |
| Radiolarians | -4.332 | 0.152 | -28.550 | $< 0.001$ |

**Table S4.** Summary table of fixed taxa novel community probability model.

| | Estimate | SE | Z-value | P-value |
|---|---|---|---|---|
| Diatoms | -3.763 | 0.140 | -26.826 | $< 0.001$ |
| Foraminifera | -3.896 | 0.105 | -36.995 | $< 0.001$ |
| Nannoplankton | -4.161 | 0.101 | -41.293 | $< 0.001$ |
| Radiolarians | -3.890 | 0.123 | -31.752 | $< 0.001$ |

#### 3.2 Transition probability model summary

**Table S5.** Summary of observed to expected transition probability comparison.

| | Observed | | | Expected | | | Observed:Expected | | |
|---|---|---|---|---|---|---|---|---|---|
| | Mean | Lower 95% CI | Upper 95% CI | Mean | Lower 95% CI | Upper 95% CI | Mean | Lower 95% CI | Upper 95% CI |
| Back $\rightarrow$ Back | 0.876 | 0.862 | 0.888 | 0.855 | 0.867 | 0.843 | 1.024 | 1.045 | 1.003 |
| Back $\rightarrow$ Instant | 0.020 | 0.017 | 0.023 | 0.036 | 0.042 | 0.029 | 0.566 | 0.716 | 0.440 |
| Back $\rightarrow$ Cumul | 0.015 | 0.013 | 0.018 | 0.016 | 0.019 | 0.014 | 0.949 | 1.178 | 0.755 |
| Back $\rightarrow$ Novel | 0.014 | 0.012 | 0.016 | 0.017 | 0.020 | 0.015 | 0.809 | 0.978 | 0.662 |
| Instant $\rightarrow$ Back | 0.026 | 0.023 | 0.030 | 0.036 | 0.042 | 0.029 | 0.753 | 0.940 | 0.594 |
| Cumul $\rightarrow$ Back | 0.013 | 0.011 | 0.016 | 0.016 | 0.019 | 0.014 | 0.811 | 1.036 | 0.625 |
| Novel $\rightarrow$ Back | 0.009 | 0.007 | 0.012 | 0.017 | 0.020 | 0.015 | 0.539 | 0.710 | 0.400 |
| Instant $\rightarrow$ Instant | 0.009 | 0.006 | 0.012 | 0.001 | 0.002 | 0.001 | 6.065 | 9.266 | 3.767 |
| Instant $\rightarrow$ Cumul | 0.001 | 0.000 | 0.002 | 0.001 | 0.001 | 0.001 | 1.164 | 2.671 | 0.402 |
| Instant $\rightarrow$ Novel | 0.002 | 0.001 | 0.003 | 0.001 | 0.001 | 0.001 | 2.483 | 4.561 | 1.204 |
| Cumul $\rightarrow$ Instant | 0.002 | 0.001 | 0.003 | 0.001 | 0.001 | 0.001 | 3.181 | 4.694 | 2.064 |
| Cumul $\rightarrow$ Cumul | 0.001 | 0.001 | 0.002 | 0.000 | 0.000 | 0.000 | 3.231 | 5.345 | 1.809 |
| Cumul $\rightarrow$ Novel | 0.001 | 0.001 | 0.002 | 0.000 | 0.000 | 0.000 | 3.540 | 5.650 | 2.076 |
| Novel $\rightarrow$ Instant | 0.007 | 0.006 | 0.009 | 0.001 | 0.001 | 0.001 | 10.506 | 13.902 | 7.760 |
| Novel $\rightarrow$ Cumul | 0.001 | 0.000 | 0.001 | 0.000 | 0.000 | 0.000 | 2.089 | 3.728 | 1.057 |
| Novel $\rightarrow$ Novel | 0.002 | 0.001 | 0.003 | 0.000 | 0.000 | 0.000 | 4.876 | 9.237 | 2.265 |

#### 3.3 Demographic model summaries

Tables S6-S11 are model summaries of binomial generalized linear mixed-effects models used to estimate differences in per-taxon probabilities of taxon loss, taxon gain, local extinction and origination, and emigration and immigration, across novel and non-novel community transitions. The fixed effects in these models are the classification of the preceding community ("PC"), the classification of the succeeding community ("SC") and the time lag between the two. The local extinction and local origination models include an additional fixed effect, the number of sampling bins to the end and start of the time-series, respectively.

**Table S6.** Model summary of overall taxonomic loss GLMM.

| Fixed effects | Estimate | Std. Error | $z$-value | $p$-value |
|---|---|---|---|---|
| (Intercept) | -0.884 | 0.066 | -13.304 | < 0.001 |
| PC-Cumul | 0.020 | 0.047 | 0.430 | 0.668 |
| PC-Instant | 0.214 | 0.035 | 6.199 | < 0.001 |
| PC-Novel | -0.015 | 0.056 | -0.273 | 0.785 |
| SC-Cumul | 1.119 | 0.037 | 30.159 | < 0.001 |
| SC-Instant | 1.277 | 0.038 | 33.689 | < 0.001 |
| SC-Novel | 1.805 | 0.048 | 37.458 | < 0.001 |
| Time lag between bins | 0.105 | 0.007 | 15.502 | < 0.001 |

| Random effects | Variance |
|---|---|
| Among planktonic taxonomic groups | 0.010 |
| Among Longhurst provinces within taxonomic groups | 0.187 |

**Table S7.** Model summary of overall taxonomic gain GLMM.

| Fixed effects | Estimate | Std. Error | $z$-value | $p$-value |
|---|---|---|---|---|
| (Intercept) | -0.874 | 0.072 | -12.131 | < 0.001 |
| PC-Cumul | 0.692 | 0.040 | 17.222 | < 0.001 |
| PC-Instant | 0.432 | 0.033 | 13.203 | < 0.001 |
| PC-Novel | 0.673 | 0.047 | 14.447 | < 0.001 |
| SC-Cumul | 0.325 | 0.045 | 7.209 | < 0.001 |
| SC-Instant | 1.362 | 0.037 | 37.310 | < 0.001 |
| SC-Novel | 1.161 | 0.054 | 21.604 | < 0.001 |
| Time lag between bins | 0.131 | 0.007 | 19.457 | < 0.001 |

| Random effects | Variance |
|---|---|
| Among planktonic taxonomic groups | 0.013 |
| Among Longhurst provinces within taxonomic groups | 0.210 |

**Table S8.** Model summary of local extinction GLMM.

| Fixed effects | Estimate | Std. Error | $z$-value | $p$-value |
|---|---|---|---|---|
| (Intercept) | -3.153 | 0.057 | -55.486 | $< 0.001$ |
| PC-Cumul | 0.115 | 0.101 | 1.138 | 0.255 |
| PC-Instant | 0.352 | 0.068 | 5.162 | $< 0.001$ |
| PC-Novel | 0.297 | 0.104 | 2.841 | 0.004 |
| SC-Cumul | 0.325 | 0.088 | 3.704 | $< 0.001$ |
| SC-Instant | 0.604 | 0.068 | 8.853 | $< 0.001$ |
| SC-Novel | 0.981 | 0.074 | 13.348 | $< 0.001$ |
| Time lag between bins | 0.332 | 0.011 | 30.113 | $< 0.001$ |
| Number of bins to time series end | -0.086 | 0.019 | -4.559 | $< 0.001$ |

| Random effects | Variance |
|---|---|
| Among planktonic taxonomic groups | 0.000 |
| Among Longhurst provinces within taxonomic groups | 0.294 |

**Table S9.** Model summary of local origination GLMM.

| Fixed effects | Estimate | Std. Error | $z$-value | $p$-value |
|---|---|---|---|---|
| (Intercept) | -3.257 | 0.049 | -66.381 | $< 0.001$ |
| PC-Cumul | 0.213 | 0.085 | 2.514 | 0.012 |
| PC-Instant | 0.261 | 0.065 | 4.019 | $< 0.001$ |
| PC-Novel | 0.509 | 0.082 | 6.212 | $< 0.001$ |
| SC-Cumul | 1.280 | 0.068 | 18.800 | $< 0.001$ |
| SC-Instant | 0.186 | 0.068 | 2.719 | 0.007 |
| SC-Novel | 1.770 | 0.068 | 26.069 | $< 0.001$ |
| Time lag between bins | 0.310 | 0.011 | 29.225 | $< 0.001$ |
| Number of bins to time series end | -0.388 | 0.020 | -19.555 | $< 0.001$ |

| Random effects | Variance |
|---|---|
| Among planktonic taxonomic groups | 0.001 |
| Among Longhurst provinces within taxonomic groups | 0.181 |

**Table S10.** Model summary of emigration GLMM.

| Fixed effects | Estimate | Std. Error | $z$-value | $p$-value |
|---|---|---|---|---|
| (Intercept) | -1.158 | 0.074 | -15.584 | $< 0.001$ |
| PC-Cumul | 0.040 | 0.051 | 0.796 | 0.426 |
| PC-Instant | 0.163 | 0.036 | 4.501 | $< 0.001$ |
| PC-Novel | -0.059 | 0.060 | -0.985 | 0.324 |
| SC-Cumul | 1.045 | 0.039 | 26.563 | $< 0.001$ |
| SC-Instant | 1.138 | 0.038 | 29.714 | $< 0.001$ |
| SC-Novel | 1.467 | 0.046 | 31.702 | $< 0.001$ |
| Time lag between bins | -0.017 | 0.007 | -2.234 | 0.025 |

| Random effects | Variance |
|---|---|
| Among planktonic taxonomic groups | 0.014 |
| Among Longhurst provinces within taxonomic groups | 0.189 |

**Table S11.** Model summary of immigration GLMM.

| Fixed effects | Estimate | Std. Error | $z$-value | $p$-value |
|---|---|---|---|---|
| (Intercept) | -1.140 | 0.070 | -16.368 | $< 0.001$ |
| PC-Cumul | 0.681 | 0.041 | 16.687 | $< 0.001$ |
| PC-Instant | 0.398 | 0.034 | 11.838 | $< 0.001$ |
| PC-Novel | 0.536 | 0.048 | 11.224 | $< 0.001$ |
| SC-Cumul | -0.149 | 0.052 | -2.861 | 0.004 |
| SC-Instant | 1.327 | 0.036 | 37.060 | $< 0.001$ |
| SC-Novel | 0.275 | 0.059 | 4.672 | $< 0.001$ |
| Time lag between bins | -0.016 | 0.007 | -2.187 | 0.029 |
| **Random effects** | | | **Variance** | |
| Among planktonic taxonomic groups | | | 0.011 | |
| Among Longhurst provinces within taxonomic groups | | | 0.206 | |

**Table S12.** Comparison of novelty in observed Neptune Sandbox data with a null simulation.

| | Background | Instantaneous novelty | Cumulative novelty | Novel community |
|---|---|---|---|---|
| Observed Neptune data | 92.50% (91.54 – 93.34%) | 3.82% (3.19 – 4.58%) | 1.74% (1.49 – 2.06%) | 1.89% (1.63 – 2.19%) |
| Null simulation | 90.88% (90.72 – 91.03%) | 2.87% (2.78 – 2.96%) | 3.88% (3.77 – 3.98%) | 2.38% (2.29 – 2.46%) |
